# High-fat diet imprints a macrophage-tumor CAMP-P2RX7 axis driving pancreatic cancer initiation, plasticity and metastasis

**DOI:** 10.64898/2026.02.10.703309

**Authors:** Ana Galván-del-Rey, Juan Carlos López-Gil, Livia Archibugi, Sonia Alcalá, Julie Earl, Michele Bevere, Antonio Agostini, Aleix Noguera-Castells, Beatriz Parejo-Alonso, Alba Royo-García, Kyra Fraser, Pian Sun, Elena Zamorano-Dominguez, Blanca Rosas-Perez, Rebeca Barrero, Silvia Jiménez-Parrado, Stephen P. Pereira, Pilar Acedo, Carmine Carbone, Claudio Luchini, Patricia Sancho, Manel Esteller, Gabriele Capurso, Bruno Sainz, Mariano Barbacid, Carmen Guerra

**Affiliations:** Experimental Oncology group, Tumor Biology Programme, Spanish National Cancer Research Center (CNIO), Madrid, Spain; Pancreato-Biliary Endoscopy and Endosonography Division, Pancreas Translational and Clinical Research Center, IRCCS Ospedale San Raffaele, Milan, Italy; Cancer Stem Cells and Fibroinflammatory Microenvironment group, Cancer department, Biomedical Research Institute (IIBm) Sols-Morreale (IIBM) CSIC-UAM, Madrid, Spain; Biomarkers and personalized approach to cancer group (BIOPAC), Area 3-Cáncer, Instituto Ramón y Cajal de Investigación Sanitaria (IRYCIS), Madrid, Spain; ARC-Net Research Centre, University of Verona, Verona, Italy and Department of Diagnostics and Public Health, Section of Pathology, University of Verona, Verona, Italy; Department of Medical and Surgical Sciences, Fondazione Policlinico Universitario “Agostino Gemelli” IRCCS, Rome, Italy; Cancer Epigenetics Group, Josep Carreras Leukaemia Research Institute (IJC), Badalona, Barcelona, Catalonia, Spain; Área Cáncer, Centro de Investigación Biomédica en Red (CIBERONC), ISCIII, Madrid, Spain; Instituto de Investigación Sanitaria Aragón (IIS Aragón), Zaragoza, Spain; Institute for Liver and Digestive Health, Royal Free Hospital Campus, University College London (UCL), NW3 2QG, London, United Kingdom; Cancer Epigenetics Group, Sant Pau Research Institute (IRSantPau), Barcelona, Catalonia, Spain; Institucio Catalana de Recerca i Estudis Avançats (ICREA), Barcelona, Catalonia, Spain; Physiological Sciences Department, School of Medicine and Health Sciences, University of Barcelona (UB), Barcelona, Catalonia, Spain; Vita-Salute San Raffaele University, Milan, Italy

## Abstract

High-fat diet (HFD) and obesity are increasingly recognized as risk factors of pancreatic ductal adenocarcinoma (PDAC), yet the mechanisms by which dietary fat contribute to oncogenic transformation remain elusive. Using an inducible acinar-specific *Kras^G12V^*/*Trp53*-loss genetically-engineered mouse model of PDAC, we established early-and late-onset protocols to assess age-dependent susceptibility to HFD. Specifically, HFD accelerated tumorigenesis with poorer prognosis in early-onset mice and, strikingly, enabled full PDAC development in late-onset adult mice otherwise resistant to oncogenic transformation. Tumors arising under HFD activated a distinct transcriptional and epigenetic state enriched in pathways or genes related to stemness, plasticity, and metastatic competence, which was maintained even in tumor-derived cell lines. Mechanistically, fatty acid-educated macrophages secreted the cathelicidin antimicrobial peptide (CAMP), activating P2X purinoceptor 7 (P2RX7) signaling in tumor cells to drive a highly plastic, immune-evasive phenotype reinforced by the expression of the peptidoglycan recognition protein 1 (PGLYRP1), further shielding tumor cells from macrophage phagocytosis. Functionally, HFD-induced tumors displayed enhanced metastatic potential independent of host context. Analysis of 164 human PDAC samples revealed that elevated body-mass index (BMI) was associated to a conserved CAMP-P2RX7-CXCR4 signature, maintained despite weight loss during disease progression. Together, these findings uncover a diet-imprinted macrophage-tumor cell circuit that promotes transformation and accelerates PDAC progression, positioning it as a therapeutic vulnerability in obesity-associated pancreatic cancer.

**Statement of significance:** High-fat diet induces pancreatic tumorigenesis in adult tissue, driving metastatic competence and a plastic state, and ultimately engages a macrophage-derived CAMP-P2RX7 circuit that reinforces immune evasion and accelerates PDAC progression.

## Introduction

High-fat diet (HFD) consumption and obesity are well-established risk factors for pancreatic ductal adenocarcinoma (PDAC) (1). As processed food intake has dramatically increased worldwide over the past decades in parallel with PDAC incidence, this association gains particular relevance in the context of a disease with a dismal 5-year survival rate of approximately 13% in the US (2). Beyond general PDAC cases, the rising incidence of “early-onset” patients (diagnosed before the age of 50) underscores the contribution of exposome-related factors to disease initiation in younger populations (3). However, traditional endogenous mouse models of PDAC, although invaluable over the past 20 years, are limited by the presence of oncogenic driver mutations from embryogenesis. In contrast, patients typically acquire these mutations during adulthood, and many harbor preneoplastic lesions that never progress to PDAC (4). Thus, the role of environmental risk factors and their ability to trigger malignant transformation requires investigation in more biologically relevant models.

Our group previously demonstrated that genetically engineered mouse models (GEMMs) harboring *Kras^G12V^* mutations and *Trp53* loss, a hallmark of advanced PDAC, in the adult pancreas were resistant to oncogenic transformation (5, 6). Only upon induction of chronic pancreatitis with cerulein did mice develop pancreatic intraepithelial neoplasias (PanINs) and PDAC lesions. Acinar cell resistance to transformation appeared reversible once inflammation was established, suggesting that inflammatory events beyond pancreatitis could also drive tumorigenesis. Indeed, *Kras*-mutant acinar cells that escape environmental clearance adopt a stem-like/quiescent state (7), and cancer stem cells (CSCs) expand during the earliest stages of tumor development once inflammation-induced metaplasia arises (8). These findings highlight that inflammatory cues fuel transformed cell plasticity and tumor initiation, yet the impact of dietary habits on this process in adulthood remains poorly defined.

Although some studies have shown that HFD accelerates tumorigenesis in embryonic *Kras*-mutated GEMMs (9, 10), mechanistic insights into how diet promotes PDAC progression and metastasis in adult models remain scarce. In this work, we evaluated the transformation capacity of adult acinar cells upon *Kras^G12V^* activation and *Trp53* loss using an inducible GEMM for PDAC that enables temporal control of oncogenic expression, allowing direct comparison between early-and late-onset protocols. We found that HFD induces systemic inflammation that drives PanINs and PDAC formation. Notably, HFD-induced tumors exhibited a distinct transcriptomic program associated with enhanced plasticity and metastatic potential. Mechanistically, fatty acid-exposed macrophages promoted inflammatory signaling through the cathelicidin antimicrobial peptide (CAMP)-P2X purinoceptor 7 (P2RX7) axis, ultimately inducing this phenotype in tumor cells and revealing a potential therapeutic opportunity to target HFD-driven PDAC.

## Materials and methods

### Mice

The *Kras^+/^*^LSLG12Vgeo^;*Elas*-*tTA*/*tetO*-*Cre* (11) and the *Trp53*^lox/lox^ (12) strains have been previously described. The *Elas-tTA/tetO-Cre* transgenes drive expression of the bacterial Cre recombinase from the *Elastase* promoter under the negative control of doxycycline (DOX) (Tet-off system). For the postnatal adult protocols, DOX (2 mg/mL, Sigma Cat no. D3447) was provided in the drinking water as a sucrose (5% w/v) solution to pregnant mothers (from the time of conception) and to their offspring until the time required to allow expression of the Cre recombinase. For the early onset adulthood protocol, mice were exposed to doxycycline until 3 weeks of age and then fed a HFD 60% (Brogaarden, Cat no. D12492I) *ad libitum* or normal chow diet until 1 year or humane endpoint. For the late onset adulthood protocol, mice were exposed to doxycycline until 8 weeks and at 12 weeks of age were fed a HFD or normal chow diet until 1 year or humane endpoint (**Figure 1A**). Both female and male mice were used for all experiments and body weight was monitored weekly. Necropsies were performed in the dissection laboratory of the CNIO Animal Facility. Mice were euthanized in a CO_2_ chamber and tissue samples (pancreas, tumor, metastases, liver, lungs, intestine, white and brown adipose tissue, spleen, kidney) were collected in 10% buffered formalin for histopathology. All animal experiments were carried out in the animal facility of the CNIO and were approved by the Ethical Committee of the CNIO and the Institute of Health Carlos III (PROEX 286.8/23). Mice were genotyped by Transnetyx (Tennessee, USA).

**Figure 1.**
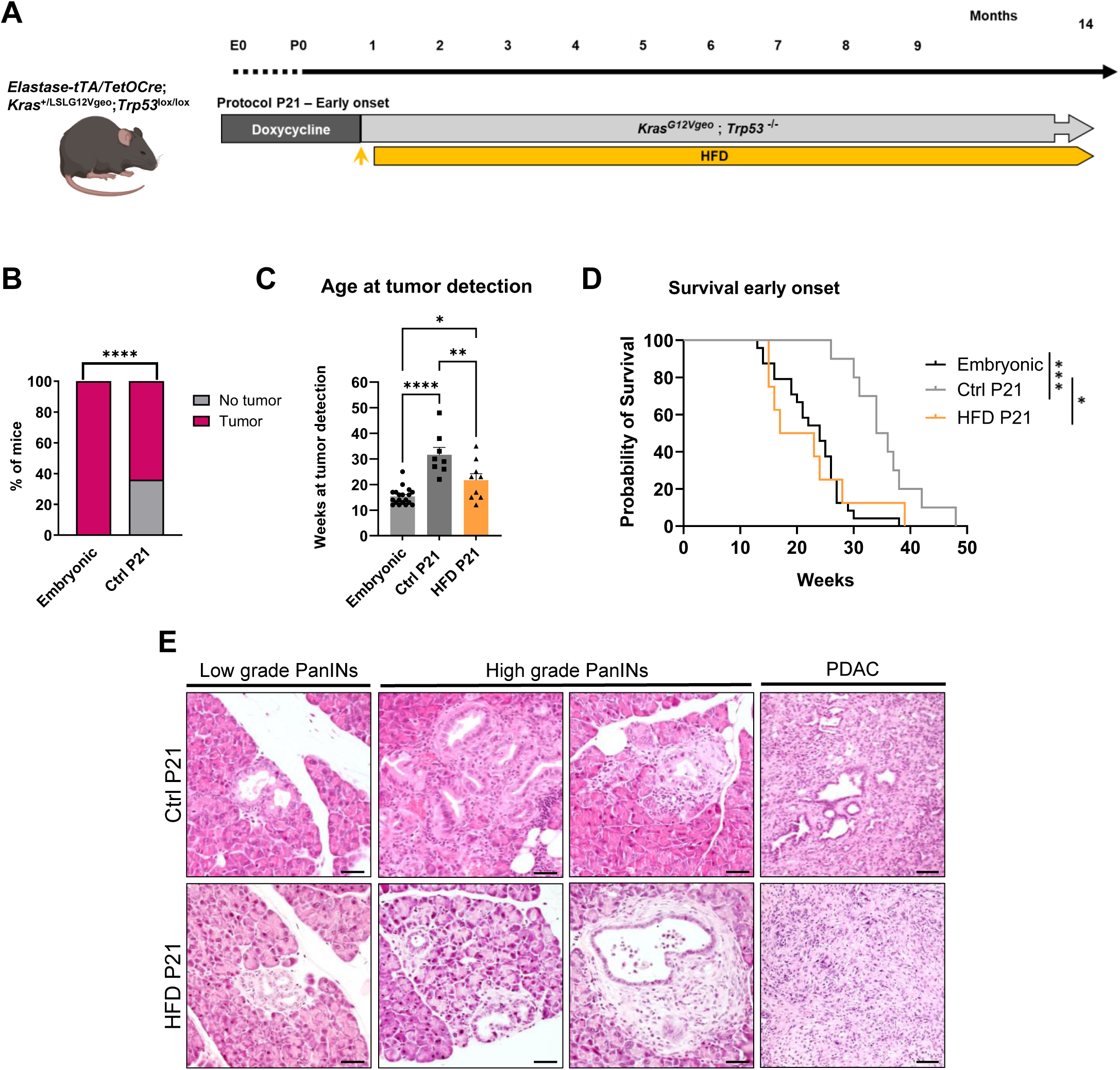
**High-fat diet promotes tumorigenesis in early-onset adult PDAC mouse models**. **A)** Representative scheme of the protocol employed to study the effect of high-fat diet (HFD) in early onset pancreatic cancer (P21). **B)** Percentage of mice from the embryonic and P21 arms in which tumors were detected (embryonic n=19, controls (Ctrl) P21 n=14). P values as determined by Chi-square test and Fisheŕs exact test. **C)** Histogram showing the weeks at tumor detection for the embryonic, Ctrl and HFD-exposed mice from the P21 protocol. Shown is the mean ± SEM (embryonic n=19, Ctrl P21 n=8, HFD n=9; P values as determined by One-way ANOVA, Tukey test). **D)** Kaplan-Meier plot showing the overall survival of mice from the embryonic and P21 (both Ctrl and HFD) settings (embryonic n=24, Ctrl P21 n=10, HFD n=8, P values as determined by Mantel-Cox test). **E)** Representative brightfield microscopy images of hematoxylin and eosin-stained pancreata at different stages of tumor development in the P21 fed with Ctrl diet (top panel) or HFD (bottom panel). Scale bar= 50 µm. P values: *, <0.05; **, <0.01; ***, <0.001; ****, <0.0001.

### Intrasplenic metastasis assay

For intrasplenic implantation, a total of 250,000 cells were resuspended in 50 μL of phosphate-buffered saline (PBS; Gibco, USA). Mice were anesthetized with 4% isoflurane (Braun Vetcare, Germany) in 100% oxygen at a flow rate of 1.5 L/min and received preoperative analgesia with buprenorphine as well as ophthalmic protection with Lacryvisc gel (Alcon, Switzerland). Body temperature was maintained using heated beds throughout the procedure. Following removal of abdominal hair, a small incision was made on the left flank to expose the spleen. The cell suspension was injected directly into the splenic parenchyma using 30-gauge insulin syringes. The peritoneal wall was closed with absorbable Vicryl sutures (Ethicon Inc., USA), and the skin was sealed with wound clips (CellPoint Scientific Inc., USA). Tumor progression was monitored weekly by abdominal ultrasound. After 21 days, mice were euthanized and tissue samples collected for the analysis of liver metastases.

### Tumor monitorization

Tumor development was assessed by weekly palpation and by ultrasound imaging performed once per month. After tumor detection, ultrasound monitoring was conducted weekly at the Molecular Imaging Core Unit of CNIO. For imaging, mice were anesthetized with 3% isoflurane (Braun Vetcare, Germany) delivered in 100% oxygen at a flow rate of 0.5 L/min. Bed heaters were used to avoid hypothermia due to the anesthesia and abdominal hair was locally removed. Tumor measurements were obtained using a high-resolution Vevo 770 ultrasound system (VisualSonics, Canada) equipped with a 40 MHz RMV704 probe (VisualSonics, Canada). Tumor sizes were calculated as (Length*Width*Height)/2.

### Primary cell lines

Murine pancreatic tumors obtained from genetically engineered mouse models were excised, segmented into smaller fragments, and enzymatically digested with collagenase P (0.2 mg/mL, Merck, Cat no. 11213857001c) and DNAse (0.5 mg/mL, Roche, Cat no. 776785) at 37°C for 15 minutes. This was followed by a 3-minute incubation at 37°C in 0.25% trypsin. The resulting cell suspensions were cultured in RPMI 1640 medium (Cat. no. R8758), supplemented with 10% fetal bovine serum and 50 units/mL penicillin-streptomycin. Epithelial clones were subsequently isolated, pooled, and expanded to establish a heterogeneous cancer cell line. Human PDAC PDX-derived cell cultures were prepared as previously described from patient-derived xenografts (21). The original PDXs were generously provided by Dr. Manuel Hidalgo through a Material Transfer Agreement with the Spanish National Cancer Centre (CNIO), Madrid, Spain (Reference no. I409181220BSMH). All primary cultures underwent routine Mycoplasma testing at intervals of no more than four weeks.

### PBMC isolation

Peripheral blood was collected in 4 mL EDTA Vacutainer tubes (BD, USA) and centrifuged at 1200xg for 10 minutes at room temperature. Plasma was then collected, aliquoted and stored at-80°C for future use. The remaining cell pellet was resuspended in 2 mL PBS, laid on top of 3 mL Ficoll-Paque (Cytiva, UK) and centrifuged at 400xg for 30 minutes with the brakes off. The PBMC buffy coat was collected, washed with 2 mL PBS mixed with 1mL red blood cell lysis buffer (Invitrogen, USA) and centrifuged at 260xg for 5 minutes. Additional washes with PBS were performed until the supernatant was clear. Patient blood samples were obtained from the Accelerated Detection of neuroEndocrine and Pancreatic TumourS (ADEPTS) biobank led by Professor Stephen Pereira. Ethical approval for this study was granted by the NHS Research Ethics Committee at Queen’s Square, London, UK (REC reference: 06/Q0512/106). Written informed consent was provided by all participants who donated samples.

### Flow cytometry

Tumor samples underwent enzymatic digestion to produce single-cell suspensions. In brief, tissues were minced using a scalpel and forceps, transferred into 50 mL conical tubes, and each tube received 5 mL of serum-free medium plus 50 μL of a digestion solution consisting of Collagenase P (0.2 mg/mL) and DNase (0.5 mg/mL). Samples were incubated at 37°C for 30 minutes under continuous agitation. To halt digestion, 1 mL RPMI supplemented with 10% FBS was added. After centrifuging at 1500 rpm for 5 minutes, supernatants were discarded. ACK buffer-mediated erythrocyte lysis was carried out by incubation at room temperature for 5 minutes. Following this, cells were washed in DMEM, centrifuged again, the supernatant removed, and resuspended in flow cytometry buffer (PBS with 2% FBS and 3 mM EDTA).

For cell lines, PBS 1X washing steps preceded trypsinization with Trypsin/EDTA 0.25% (Lonza, Cat no. CC-5012). Post-trypsinization, the cells were resuspended in complete RPMI medium. When analyzing murine cells, cell suspensions were treated with purified anti-mouse CD16/CD32 antibodies (eBioscience, Cat no. 14-9161-73) to block Fc receptor binding. Human cell preparations were incubated for 15 minutes at 4°C with blocking buffer (PBS with 5% FBS and 3 mM EDTA) to reduce non-specific binding.

Antibodies used for surface marker staining are detailed in **Supplementary Table 1**. For excluding non-viable cells, samples were stained with DAPI at 2 µg/mL (4’,6-diamidino-2-phenylindole; Sigma, Cat no. D9542-5MG). Cells were then suspended in flow cytometry buffer [1X PBS, 3 mM EDTA (v/v)] prior to analysis. Flow cytometry was performed on an Attune™ NxT Acoustic Focusing Cytometer equipped with four lasers (Thermo Fisher Scientific), and data analysis was completed using FlowJo software (Tree Star Inc., Ashland, OR).

### Histopathology

For histopathological analyses, tissues were serially sectioned (3 μm thick), and every 10th section was stained with hematoxylin and eosin (H&E) by the CNIO Histopathology unit. Representative sections were chosen for the grading and enumeration of lesions and quantification of tissue damage. For H&E-stained slides of pancreata, areas were marked as normal tissue, ADM, low grade and high grade PanINs or PDAC tumor tissue. Moreover, tumors were classified as differentiated or poorly differentiated using the guidelines from the College of American Pathologists (https://documents.cap.org/protocols/cp-pancreas-exocrine-17protocol-4001.pdf).

### Immunohistochemistry

Specimens were fixed in 10% formalin (Sigma) and embedded in paraffin. Immunohistochemistry staining was performed by the Histopathology Unit of CNIO. For histopathological analysis, specimens were serially sectioned (3 μm thick) and stained with H&E. Antibodies used for mouse tissue are included in **Supplementary Table 1.** Images were analyzed and exported using QuPath software (v0.5.0). For human tissue, patients have been selected from a cohort already published (13). Immunohistochemistry has been performed as detailed before (14). Antibodies used for human samples are included in **Supplementary Table 1**. For evaluating the expression, a combined semiquantitative score was adopted, taking into account both the percentage score, which is obtained based on the percentage of stained cells (score 0 = 0%, 1 = 1-25%, 2 = 26-50%, 3 = 51-75%, 4 = 76-100%) and the intensity score, which is based on the intensity of the staining (score 0 = no stained cells, 1 = weak staining, 2 = moderate staining, 3 = strong staining). For obtaining the value of combined semiquantitative score, the value of percentage score and the value of intensity score are multiplied. Thus, the combined / semiquantitative score ranges from 0 to 12. The evaluation of the scores has been performed on neoplastic cells and immune cells.

### Immunofluorescence and microscopy

Microscopy brightfield images were captured with an EVOS FL Microscope (Invitrogen, Cat no. AMF4300). For immunofluorescence, FFPE sections were prepared at the CNIO Histopathology unit, deparaffinized in graded ethanol, and underwent antigen retrieval using 1 mM EDTA. After blocking (PBS 1x, 1% BSA, 2% FBS), primary antibodies were incubated overnight at 4°C, followed by 1 hour at 37°C with secondary antibodies and DAPI. Samples were mounted in ProLong (Invitrogen, Cat no. P36970) and imaged with a Confocal Stellaris 8 STED microscope (Leica). Image analysis was performed using ImageJ.

### RNA extraction and RT-qPCR

For qPCR analysis, RNA was extracted using guanidine thiocyanate as described before (15) and cDNA synthesized with the SS II cDNA synthesis kit (Cat no. 18064022, Invitrogen) followed by RT-qPCR analysis using SYBR green and a StepOne Plus real-time thermo-cycler (Applied Biosystems). The thermal cycle consisted of an initial denaturation step of 10 min at 95°C followed by 40 cycles of denaturation (15 s at 95°C) and annealing (1 min at 60°C). The results obtained for each gene were normalized with the β-actin levels. Primers are listed in **Supplementary Table 2**.

### Library preparation for RNA-seq

RNA concentration was determined using a NanoDrop spectrophotometer, and RNA quality was evaluated based on the RNA Integrity Number (RIN) using the LabChip GX Touch Nucleic Acid Analyzer (PerkinElmer). RNA quality control, library preparation, and sequencing were performed at the CNIO Genomics Unit. For library generation, 1 μg of total RNA was processed using the NEBNext Ultra II Directional RNA Library Prep Kit for Illumina (NEB #E7760), following the manufacturer’s recommendations. Briefly, the poly(A)+ RNA fraction was isolated and fragmented, followed by synthesis of double-stranded cDNA. Subsequent enzymatic steps included end-repair, dA-tailing, and adapter ligation. The adapter-ligated products were amplified by PCR using Illumina paired-end primers. Final libraries were loaded onto an Illumina flow cell for cluster generation and sequenced on an Illumina NextSeq 500 system according to the manufacturer’s protocol.

### DNA extraction and Illumina Mouse Methylation microarrays

Genomic DNA was extracted at the Genomics Unit of the Josep Carreras Leukemia Research Institute. Briefly, DNA was isolated from cell pellets using the DNeasy Blood & Tissue Kit (Qiagen, Hilden, Germany) and quantified with a Qubit fluorometer. DNA integrity was assessed by agarose gel electrophoresis, and samples (≥600 ng per sample) were bisulfite-converted using the EZ-96 DNA Methylation Kit (Zymo Research Corp., CA, USA) according to the manufacturer’s instructions. Bisulfite-converted DNA was then hybridized to Infinium Mouse Methylation microarrays as previously described by García-Prieto *et al.* (16).

### Endoscopic ultrasound database generation

Upon approval from the Internal Review Board (IRB BIO-PANCREAS, version 3, 2021), specimens were collected from patients diagnosed with pancreatic ductal adenocarcinoma (PDAC) at San Raffaele Research Hospital. Eligible subjects included treatment-naïve individuals with clinical histories and radiological findings indicative of PDAC. Patients were not involved in the design, implementation, analysis, or dissemination of the study. Informed consent was obtained prior to conducting diagnostic EUS procedures under deep intravenous sedation with Propofol (Diprivan, Zeneca, Germany). Procedures utilized a Pentax therapeutic linear echoendoscope (EG3870UTK, EG38J10UT) in conjunction with Hitachi ultrasound systems (Arietta 850, Arietta V70). EUS-guided tissue acquisition (TA) was performed using a 25G fine-needle aspiration (FNA) needle (Expect SlimLine, Boston Scientific) employing the slow-pull technique. RNA was subsequently isolated, quantified, and evaluated according to established protocols (17).

### RNA-sequencing analysis

FASTQ file integrity was verified using MD5 checksum comparison, and read quality was assessed with FastQC, with optional contamination screening via FastQ Screen; all quality metrics were summarized in a MultiQC report. Reads from different lanes were concatenated, optionally downsampled to a fixed number of reads using a defined seed and trimmed for adapters and low-quality bases using bbduk (BBTools). Reads were then aligned using STAR, HISAT2, or Salmon as specified in the configuration; STAR and HISAT2 used genome indices generated from a reference FASTA and GTF annotation (with HISAT2 outputs subsequently sorted using samtools), while Salmon performed pseudo-alignment using a transcriptome-based index. For STAR and HISAT2 alignments, gene-level quantification was performed with htseq-count or featureCounts after indexing BAM files with samtools, using the same GTF annotation employed for alignment. Per-sample count files were merged into a final counts matrix for downstream differential expression analysis. Differentially expressed genes (DEGs) were identified from count matrices using DESeq2 in R, excluding genes with fewer than five counts. The Wald test was employed to determine significant changes in expression between conditions, while Principal Component Analysis (PCA) was used to evaluate variability among samples. DEGs for each condition were selected considering Log2FC ≠ 0 and p ≤0.05 and subsequently ranked according to their Log2FC value. Gene set enrichment analysis (GSEA) was performed using annotations from Reactome, KEGG, and MSigDB (www.broadinstitute.org/gsea/index.jsp), as well as custom gene sets described in **Supplementary Table 3**. In addition, genes were ranked using moderated t statistics from limma. After applying the Kolmogorov-Smirnov test, gene sets with FDR < 0.25 were considered significantly enriched.

For the EUS dataset, we performed two kinds of analyses. For the differentially expressed genes in patients with body mass index (BMI) <25 or ≥25, we employed limma package (R/Bioconductor). A design matrix was constructed including BMI group as the primary variable of interest and adjusting for relevant covariates (sex, age, smoking status, batch, and RNA extraction methodology). Linear models were fitted to the expression matrix using lmFit, followed by empirical Bayes moderation with eBayes. Differentially expressed genes for the *Obese vs. reference BMI group* contrast were extracted using topTable. To assess global transcriptional patterns and sample structure, a principal component analysis (PCA) was conducted, and samples were visualized according to BMI group. Differential expression results were further summarized using a volcano plot highlighting genes with nominal *p* < 0.05. Significant genes were filtered based on nominal significance and exported with their log fold changes, *p*-values, and adjusted *p*-values for downstream interpretation. For a set of predefined stem-related genes, we generated violin plots to visualize their expression distribution across BMI groups. For each gene present in the expression matrix, a sample-level dataset was constructed by pairing log-RPKM expression values with the corresponding BMI group. Violin plots were generated using ggplot2, combining kernel density estimates with overlaid boxplots to summarize central tendency and dispersion. Group differences in gene expression were statistically evaluated using the Wilcoxon rank-sum test, and the resulting *p*-values were annotated directly on each plot.

For the organoid implantation assay dataset (GSE148135), Raw counts were normalized and converted in logCPM using edgeR (R package). For the figure, the p values refer to Dunn_test that was used post Kruskall-Wallis to test difference in expression.

For the PBMC RNA-seq, RNA was extracted from isolated PBMC samples using the RNeasy Mini Kit (Qiagen, Germany), according to the manufacturer’s recommendations. RNA concentration and integrity were measured using a NanoDrop™ 2000 Spectrophotometer (ThermoFisher, UK) and TapeStation RNA High sensitivity assay (Agilent, USA), respectively. In collaboration with UCL Genomics (UCLG), cDNA libraries were prepared using the Watchmaker RNA Library Prep Kit with Polaris rRNA and Globin Depletion (Watchmaker Genomics, USA). Libraries were indexed using IDT xGen unique dual index (UDI) adapters incorporating unique molecular identifiers (UMIs) and sequenced with the NovaSeq 6000 (S4 v1.5 flow cell, 300 cycles, XP workflow), to generate paired-end 150 bp reads, yielding approximately 40 million read pairs per sample. Demultiplexing, quality control (FastQC and MultiQC), and transcript quantification using Salmon (pseudo-mapping) were performed by UCLG to generate gene-level raw count matrices for downstream analysis. Data normalization and differential gene expression analysis were conducted using DESeq2 via the SARTools package, developed by Hugo Varet *et al.* (17), and analysis quality was assessed using principal component analysis (PCA) and additional diagnostic visualizations. Significantly differentially expressed genes were identified as those with an adjusted p value <0.05 and log2 fold-change<1 between comparison groups.

### DNA methylation analysis

Raw signal intensity files (IDATs) were processed using standard settings in the SeSAMe R package. Unsupervised analyses of the global methylome were performed using t-SNE. Supervised analyses were conducted by calculating mean methylation values per condition and per CpG site, and CpG sites with absolute methylation differences between conditions ≥0.33 were considered biologically meaningful. Differential methylation was assessed using a linear regression model implemented in the limma R package, and CpG sites with an adjusted P-value ≤0.05 were considered significant. CpG sites were annotated using the annotation described in García-Prieto *et al*., and downstream analyses included gene set enrichment analysis of gene-associated CpG sites using the EnrichR R package and the Panther database. Pathways were considered significant if they had adjusted P-values ≤0.05.

### hCAP-18/LL-37 ELISA

Patients with suspected or confirmed pancreatic ductal adenocarcinoma (PDAC) were recruited from the General Surgery, Gastroenterology, and Medical Oncology departments of Ramón y Cajal University Hospital (Madrid, Spain). The study was approved by the local ethics committee, and all participants provided written informed consent. Blood samples were collected at study entry, prior to any oncological treatment, for serum isolation. Serum samples from healthy individuals without a history of digestive disease or cancer were obtained from the Ramón y Cajal-IRYCIS Biobank (PT20/0045; PT23/00098), part of the Spanish National Biobanks Network (ISCIII Biobank Register No. B.0000678). Serum LL-37 levels were quantified using a commercial ELISA kit (ELabScience, E-EL-H2438) according to the manufacturer’s instructions. Samples were diluted 1:10 and incubated on plates pre-coated with LL-37-specific capture antibodies. Following incubation with detection antibodies and chromogenic substrate, absorbance was measured at 450 nm with correction at 540 nm. Concentrations were calculated by interpolation from a standard curve and expressed as ng/mL. All samples were analyzed in duplicate, and those with intra-assay coefficients of variation >15% were excluded.

### Macrophage isolation

Murine bone marrow-derived cells (BMDCs) were harvested from the femurs and tibiae of a minimum of three mice using centrifugation. The resulting BMDC pellets were resuspended in RPMI 1640 medium (Invitrogen, Cat. No. 61870044) and incubated for 24 h to facilitate monocyte attachment. Subsequently, monocytes were differentiated into macrophages under adherent conditions for 5-7 days on non-tissue culture-treated 100 mm dishes in RPMI supplemented with 10% FBS and 10 ng/mL murine M-CSF (PeproTech, Cat. No. 315-02). Additionally, certain experiments utilized immortalized murine bone marrow-derived macrophages generously provided by Dr. Antonio Castrillo.

### Macrophage conditioning and co-culture assays

After differentiation, for macrophage polarization, they were cultured without stimulation to obtain untrained conditioned medium or with stimulation (fatty acids) for 24 h to obtain a lipid loaded state. Then, polarization medium was removed and RPMI was left to be conditioned for 48 h to obtain fatty acid trained-macrophage conditioned medium.

### Phagocytosis assay

Five days after isolation of macrophages, tumor fluorescent cells were harvested and seeded together with macrophages during 24 h. Afterwards, cells were detached using scrapers and macrophages phagocytosing fluorescent tumor cells were measured by flow cytometry, as described above, gating on the double-positive population: F4/80 and reporter genes (i.e., YFP).

### Stimulation assays

For stimulation assays, tumor or immune cells were seeded in 6-well plates and treated with 1 mL of complete RPMI medium plus recombinant CAMP (5 µg/mL, Cat. no. HY-P4855, MedChemExpress) during 24h/48h, fatty acids (50 µM, oleic acid; 50 µM sodium palmitate; 200 µM linoleic acid; Cat no. O3008, P9767, L9530) or MØ-derived conditioned media. After indicated time points, cells were trypsinized or scraped and harvested for flow cytometry of the indicated markers or RNA-seq analysis.

### Spheroid formation assay

Cancer stem cell-enriched spheroids were cultured employing 10,000 cells per well in ultra-low attachment plates (Corning) during 4-7 days using serum-free DMEM/F12 medium (Invitrogen, Cat no. 21331046) with B27 (1:50; Invitrogen, Cat no. 17504044), 20 ng/mL bFGF (PAN-Biotech, Sigma, Cat no. GF446-10UG), L-Glutamine (Invitrogen, Cat no. 25030081) and 50 units/mL penicillin-streptomycin (Invitrogen, Cat no. 11548876).

### Transwell invasion assay

Cell invasion assays were performed using 24-well Transwell chambers (Corning, NY) equipped with polycarbonate inserts containing 8 μm pores. Inserts were coated with 50 μL of Matrigel (Corning, NY) diluted 1:3 in culture medium. A total of 50,000 cells suspended in serum-free medium were seeded into the upper chambers, while the lower wells were filled with medium containing 1% FBS as a chemoattractant. After 24 h of incubation, non-invading cells on the upper surface of the membrane were gently removed with a cotton swab. Cells that had migrated to the lower membrane surface were fixed with 0.1% glutaraldehyde, stained with 0.5% crystal violet, and visualized under a light microscope. The number of invaded cells was quantified by counting five randomLy selected microscopic fields.

### Wound healing assay

Cells were seeded (100,000) into 24-well plates and grown until reaching approximately 95% of confluent monolayers. A straight line was scratched through the monolayer with a 10 μL pipette tip to create a wound. Then, the closure of the wound was visualized using an EVOS phase-contrast microscopy and the gap distance between the edges was calculated using ImageJ software.

### Statistical analysis

Results are reported as means ± standard error of the mean (SEM) or standard deviation (STDEV), as specified. Differences between group means were assessed using Student’s t-test, unless otherwise indicated in the figure legends. P values less than 0.05 were deemed statistically significant, while non-significant outcomes are denoted as “ns”. All statistical analyses were conducted using GraphPad Prism version 8.0 (San Diego, California, USA).

## DATA AVAILABILITY

### Materials availability

All data related to the results are available as part of the article. Additional information will be available on request.

### Data and code availability

Transcriptional and methylation data generated in this study have been deposited in NCBI-SRA BioProject ID PRJNA1422783 (HFD RNA-seq) and in GEO ID GSE319337 (methylation data). Unique identifiers for publicly available datasets are indicated, a list of figures that have associated raw data can be provided, and there are no restrictions on data availability. Data from the EUS dataset can be provided upon reasonable request and DTA signing with the San Raffaele Hospital. This paper does not report any original code. Further information if required is available from the lead contact upon request.

## Results

### HFD promotes pancreatic tumorigenic transformation in an early adulthood onset model of PDAC

To study the contribution of high-fat diet in pancreatic tumorigenesis we took advantage of an inducible GEMM of PDAC with KRAS^G12V/+^ and P53^KO^, which mimics the mutational characteristics of advanced PDAC (*Elas*-tTA/*tetO*-Cre; *Kras^LSL^*^G12V/+^; *Trp53*^lox/lox^). We developed two risk factors exposure protocols to model the early and late onset development of the disease, in addition to the classical embryonic setting as a reference. For the early onset (P21), we maintained mice on doxycycline (DOX) for three weeks and then DOX was removed and exposure to high-fat diet (HFD) was initiated. As DOX requires approximately one month for complete clearance, HFD exposure prior to full induction of the mutations repressed by DOX resembles adolescents with HFD-like habits before mutation acquisition (**Figure 1A**). Mice were kept on HFD until one year of age or humane endpoint. For the embryonic setting, *Elas*-tTA/*tetO*-Cre; *Kras^LSL^*^G12V/+^; *Trp53*^lox/lox^ mice were kept without DOX allowing for transgenes recombination during embryonic development leading to 100% disease development. As expected, both female and male mice gained weight after HFD-exposure (**Figure S1A**) and the expected systemic effects caused by the diet were assessed, including fatty liver (**Figure S1B**), acinar-to-adipose replacement in the pancreas, intra-acinar lipid droplets accumulation (**Figure S1C**), and brown to white adipose tissue transformation (**Figure S1D**). We further confirmed that HFD exposure alone does not lead to preneoplastic lesions (PanINs) formation in wild-type (WT) mice, demonstrating that the diet by itself does not trigger neoplastic transformation (**Figure S1E**).

As reported in previous studies from the laboratory, adult acinar cells present an oncogenic transformation resistance that increases with age, with no tumor formation when mutations are induced at 60 days post-birth (5, 6) (**Figure 1B**). Interestingly, P21 mice exhibited an intermediate phenotype, showing preneoplastic lesions and full PDAC, but with an increased resistance to tumor development compared with the embryonic model (**Figure 1B**). Notably, P21 mice exposed to HFD developed tumors earlier than P21 control mice, although later than the embryonic model (**Figure 1C-D**). Despite these differences, survival was comparable between the P21 HFD protocol and the embryonic model, and in both cases significantly shorter than in the P21 control group (**Figure 1D**). Histological analysis of these tumors revealed distinct tumor phenotypes, with control tumors displaying a more differentiated epithelial morphology characterized by ductal-like structures, whereas tumors arising under HFD conditions mainly exhibited a squamous/mesenchymal-like phenotype (**Figure 1E**). These results suggest that HFD promotes early onset of PDAC tumors, affecting survival in an early adulthood context.

### HFD-promoted pancreatic tumors displayed a mesenchymal-like phenotype

Based on the histological differences observed before, we conducted a marker-based characterization of the P21 control and HFD-promoted tumors. We found that control tumors displayed high levels of cytokeratin 19, a common marker of epithelial tumors, while HFD-promoted tumors expressed markedly lower levels (**Figure 2A**). On the other hand, vimentin, a marker of mesenchymal tumors, was overexpressed in HFD tumor cells, while in control tumors it was mainly restricted to fibroblasts (**Figure 2A**). Consistently, the epithelial marker EpCAM, was also highly reduced in HFD-promoted tumors (**Figure 2B-C**). Interestingly, in the HFD-promoted tumors we observed a population of cells that expressed both EpCAM and vimentin, suggesting that HFD could induce a hybrid epithelial/mesenchymal phenotype (**Figure 2B-C**).

**Figure 2.**
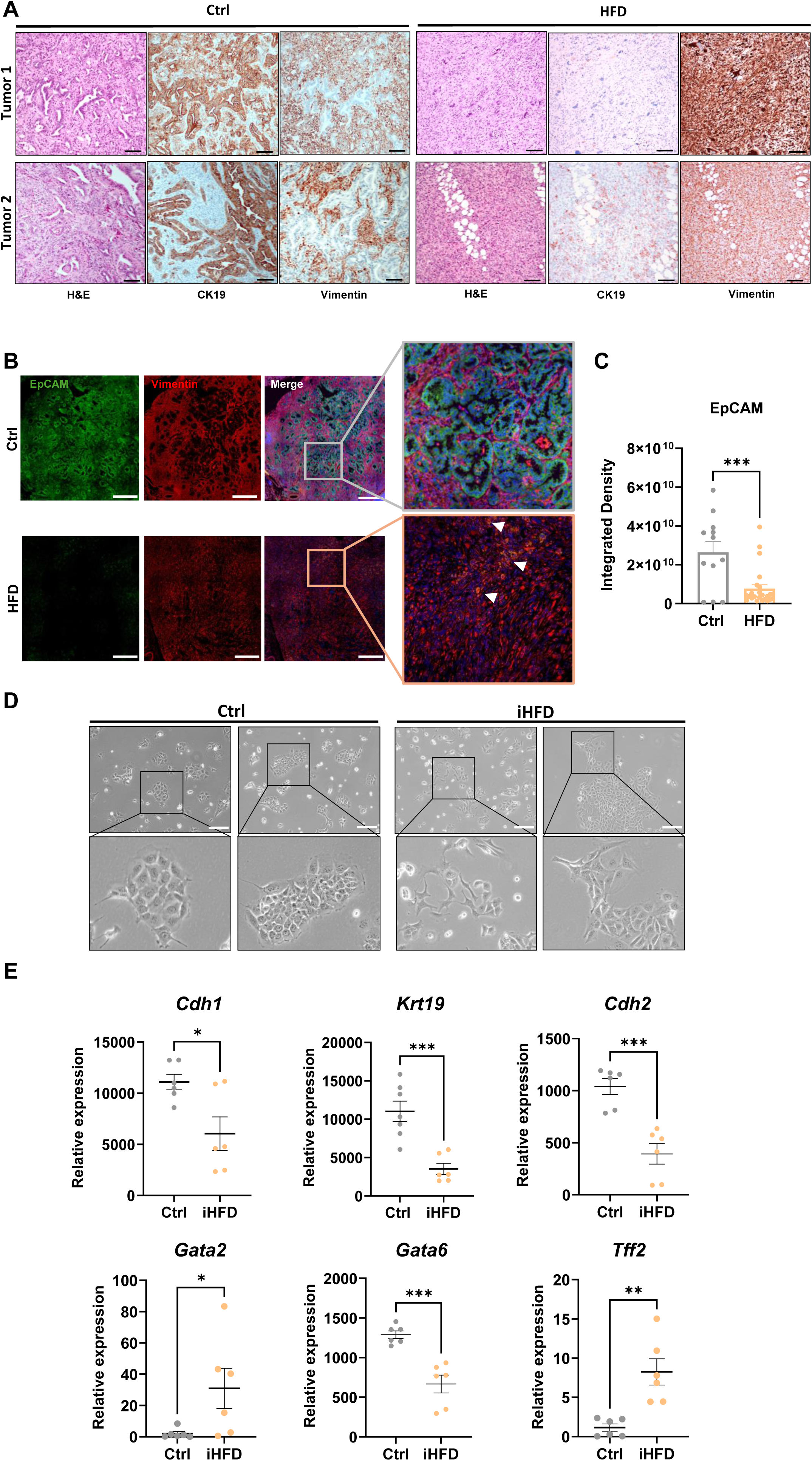
Histology and morphological characterization of control and high-fat diet-promoted tumors and primary tumor-derived cells. A) Representative brightfield microscopy images of hematoxylin and eosin (H&E), along with cytokeratin 19 (CK19) and vimentin immunohistochemistry from two tumors generated in P21 control (Ctrl) and HFD-fed mice. Scale bar= 50 µm. B) Representative IF confocal microscopy images from Ctrl and HFD tumors stained with EpCAM (green, epithelial tumor cells), Vimentin (red, mesenchymal tumor cells and fibroblasts) and DAPI (blue, nuclei). Arrowheads indicate double-positive EpCAM/Vimentin tumor cells. Scale bar= 500 µm. C) Quantification of the integrated density for EpCAM staining in Ctrl and HFD tumors. Shown is the mean ± SEM for each measured field (n=7 mice, 3 for Ctrl and 4 for HFD; P values as determined by unpaired T-test). D) Representative brightfield microscopy images of P21 Ctrl and HFD-induced (iHFD) primary tumor-derived cell lines. Scale=50µm. E) Relative expression of mRNA levels measured by RT-qPCR of the indicated genes in Ctrl and iHFD primary tumor cell lines. Shown is the mean ± SEM (n=6, P values as determined by unpaired T-test). P values: *, <0.05; **, <0.01; ***, <0.001; ****, <0.0001.

To better understand the molecular changes produced by HFD in tumor cells, we derived primary cell cultures from both P21 control and HFD-induced tumors. As expected, and consistent with the phenotype observed *in vivo*, control cell lines exhibited an epithelial morphology in culture, whereas HFD-induced (iHFD) tumor cell lines displayed a mesenchymal morphology (**Figure 2D**). In line with this, iHFD cell lines showed a reduction of *Krt19, Cdh1 and Gata6*, genes associated with an epithelial/classical-like phenotype. Likewise, we found an upregulation of *Gata2*, which regulates *Gata6* expression in an opposite manner (18, 19). Unexpectedly, we observed a decrease in *Cdh2* and an increase in *Tff2,* mesenchymal and epithelial-associated markers respectively (**Figure 2E**). These results indicated that HFD induced a hybrid phenotype between classical/epithelial and basal/mesenchymal states and suggested that HFD promotes a plastic state in tumor cells.

### HFD induces transcriptomic and epigenetic changes in tumor cell state

To interrogate in-depth the transcriptional state of iHFD tumor cells, we performed RNA sequencing analysis of six control and six iHFD P21 cell lines. We first found that iHFD cells exhibited higher transcriptomic differences among themselves than control cells (**Figure 3A**). We next performed a differential gene expression analysis, and found significantly upregulated genes associated with extracellular matrix remodeling (i.e., *Fgfr1, Loxl1, Mmp7, Has2*), stemness (i.e., *Prom2, Pglyrp1, Isg15, Agr2, Fn1*), mesenchymal identity and metastatic behavior (i.e., *Snai1, Zeb2, Grem1, Ngfr, Plau, Foxd1, Foxc1*), and immune evasion (i.e., *Csf1r, Cxcl5, Thbs1, Ccl7, Ccl9, Pglyrp1*) (**Figure 3B**). Among the downregulated genes in iHFD we found mainly genes associated with mitochondria activity (*mt-Co3, mt-Nd3, Ndufb4*). In addition, we found that *Gata2* was upregulated, consistent with the aforementioned data, sustaining the idea of the extinction of the classical subtype program (**Figure 3B**). To better characterize these transcriptomic changes, we performed Gene Set Enrichment Analysis (GSEA) employing Molecular Signature Database and custom gene sets. Interestingly, we found upregulated a gene signature related to increased tumorigenesis by chemical induction (**Figure 3C)**, as expected from a tumor induced by an external agent such as diet. Apart from this, we found pathways mainly associated to epithelial-to-mesenchymal transition (EMT) and the acquisition of mesenchymal identity and trajectory (20), stemness and plasticity, inflammation and immune system modulation (IL1β and interferon response, and antimicrobial response), ECM organization (FGF-response and collagen organization), and, as expected under HFD-induced nutrient overload, nutrient response (**Figure 3C, S2A-E**). Based on this, we also compared the expression of marker genes associated with plasticity and stemness in PDAC (8, 21–33), and we found that iHFD cells presented a general increase in the expression of some (but not all) genes including *P2rx7, Pglyrp1, Agr2, Sox9,* and *Ror2* (**Figure 3D**). As this hybrid/plastic phenotype remains stable in the iHFD cell lines, we evaluated epigenomic changes at the DNA methylation level for these cell lines. Interestingly, this analysis suggested that the global methylome clusters the samples per group (**Figure S2F**). We calculated the mean methylation value per CpG sites per group, and the difference among groups. We selected the CpG sites that had an absolute methylation difference higher than 0.33 between groups, and then, we applied a linear regression model, obtaining 2,526 CpG sites differentially methylated (adjusted p-value<0.05). From them, 92.12% (n=2,327) were hypomethylated in iHFD in comparison to the control group (**Figure S2G**), suggesting that HFD is likely to promote gene upregulation. Moreover, most of these CpG sites that were differentially methylated were located at the 5’ regulatory region of the genes (42.91%; n=1,084) (**Figure S2H**). Then, we used the differentially methylated CpG sites to cluster the samples (**Figure S2I**) and we performed an enrichment analysis with EnrichR using the genes that had at least one CpG site differentially methylated. Using the Panther database, we found 2 differentially regulated pathways, the WNT signaling pathway and the cadherin pathway (**Figure S2J**), supporting that HFD influences the plastic-like EMT state of tumor cells.

**Figure 3.**
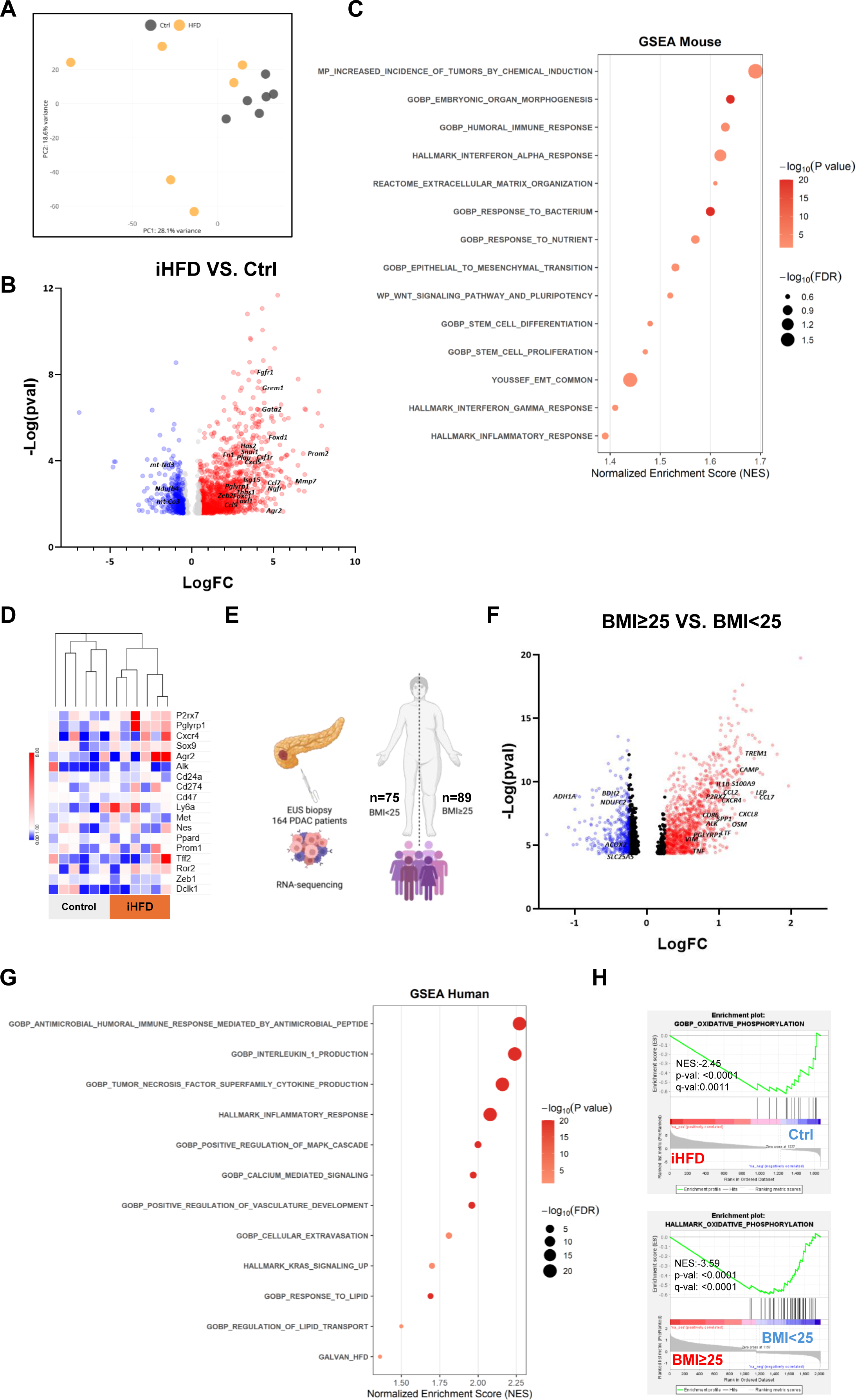
Transcriptomic profile of high-fat diet-induced cell lines and PDAC patients with high body mass index (BMI). **A)** Principal Component Analysis (PCA) for P21 control (Ctrl) and high-fat diet-induced (iHFD) primary tumor cell lines analyzed by RNA-seq. **B)** Volcano plot illustrating genes significantly up-(red) and downregulated (blue) in iHFD cell lines. **C)** Gene Set Enrichment Analysis (GSEA) plot showing the enrichment in the indicated pathways. **D)** Heat map showing the levels of expression as fold change of normalized counts for the indicated genes in Ctrl and HFD cell lines included in the RNA-seq analyses. The indicated genes are linked to stemness, plasticity and immune evasive phenotypes. **E)** Schematic representation of a patient comparison study involving 164 endoscopy-guided ultrasound (EUS) biopsy tumor samples, correlating BMI with tumor transcriptomic profiles. BMI cutoff has been set as 25 (n for BMI<25=75, n for BMI≥25=89). **F)** Volcano plot showing the genes significantly up-(red) and downregulated (blue) in patients with a BMI higher or equal to 25. **G)** GSEA plot showing the enrichment of the indicated pathways. **H)** GSEA plots depicting the enrichment of the oxidative phosphorylation pathway in both murine (top) and human (bottom) RNA-seq datasets.

To cross validate the transcriptomic results with human data, we took advantage of a database including 164 endoscopy-guided ultrasound (EUS) biopsies of PDAC patients of all stages sequenced by bulk RNA-seq (34). First, we divided patients according to their BMI at the time of diagnosis, considering equal to or higher than 25, patients with overweight or obesity; and patients with less than 25 as normal weight or lean (**Figure 3E**). When we compared both groups and performed differential gene expression analysis, we found upregulated several genes associated with inflammation (i.e., *IL1B, TNF, OSM*), antimicrobial response associated with stemness (i.e., *CAMP, PGLYRP1, LTF*), CSCs markers related also to metastases (i.e., *P2RX7, ALK, CXCR4*) and genes involved in immune evasion (*CXCL8, SPP1, S100A9, CD86, CCL7, CCL2*) (**Figure 3F**); some of them also upregulated in our murine dataset. Interestingly, GSEA analysis revealed a significant enrichment in pathways related to macrophages, including those associated with lipids, as well as inflammatory responses mediated by IL1β, TNFα, IL8, or interferons. We also observed enhanced antimicrobial activity via antimicrobial peptides (**Figure 3F, S3A**); as well as several pathways associated with oncogenic KRAS activity, ERK1/2 and MAPK signaling pathways (**Figure S3B**). Likewise, patients with higher BMI presented an enrichment in genes associated with increased blood vessel formation and enhanced migration (**Figure S3C**), as well as response to lipids (**Figure 3G**). Additionally, we generated a gene signature from our murine dataset based on the upregulated genes in iHFD cells (GALVAN_HFD) and assessed its enrichment in the human cohort. As expected, patients with high BMI were significantly enriched for this murine-derived gene signature (**Figure 3G**). Notably, both mouse and human datasets mainly showed downregulation of pathways associated with oxidative phosphorylation (**Figure 3H**). As most of these pathways were common between our human and murine datasets, we compared the genes driving these processes and found 79 commonly upregulated and 36 downregulated (**Figure S4A-B**). Remarkably, we identified P2RX7, an ATP-gated calcium channel and known CSC marker, among the common upregulated genes. P2RX7 is associated to EMT and stemness promotion acting as a receptor for the cathelicidin antimicrobial peptide (*Camp*/CAMP in mice, *CAMP*/hCAP-18/LL-37 in humans), also upregulated in patients with high BMI (**Figure 3F**) (26). These data, along with the enrichment in gene signatures associated to antimicrobial peptides and calcium signaling, led us to hypothesize that HFD could be driving the P2RX7-CAMP axis, promoting a highly plastic, stem-like state in tumor cells driving aggressiveness and tumor progression.

### HFD-induced tumor cell lines present enhanced plasticity and stemness characterized by P2RX7 overexpression

To test our hypothesis regarding enriched stemness, we cultured control and iHFD cell lines as CSC-enriched spheroids. Plating an equal number of cells, iHFD cell lines formed significantly more spheroids, supporting the plastic/stem-like nature of iHFD tumor cells (**Figure 4A-B**). Next, we evaluated the expression of P2RX7 in our control and iHFD cells by RNA-seq data, qPCR and flow cytometry. P2RX7 was found consistently and significantly overexpressed in iHFD cells (**Figure 4C-F**). Moreover, we found overexpression of P2RX7 in orthotopic tumors generated in HFD-fed C57Bl/6 mice (**Figure 4G**). Given that diet-induced alterations may partially recapitulate the molecular effects of obesity, we analyzed *P2rx7* expression in an RNA-seq dataset of murine orthotopic PDAC organoids grown in HFD-fed and in leptin-deficient (ob/ob) genetic obese mice (35). Interestingly, in both models there was a significant and consistent enrichment in *P2rx7* expression compared to controls (**Figure S5A-B**), suggesting that our HFD-fed model could mimic what occurs in patients with obesity. Thus, aiming to translate these results to patients, we evaluated the expression of *P2RX7* in our cohort at the time of PDAC diagnosis, comparing patients with high and low BMI one year prior to diagnosis and at the time of diagnosis (**Figure S6A**). We found that patients with high BMI one year before PDAC diagnosis exhibited a significant enrichment of *P2RX7* expression (**Figure 4H**), but this difference disappeared at the time of the diagnosis (**Figure 4I**). This effect could be likely due to the cachexia developed during the course of the disease. Therefore, we evaluated those patients that at the time of the diagnosis were obese but lost weight, versus those that were always lean. Interestingly, those patients with high BMI one year before diagnosis retained elevated levels of *P2RX7* expression at the time of diagnosis (**Figure 4J**), even though they were “lean” at that point. This finding suggests that P2RX7 may serve as a risk factor marker linking high BMI to PDAC.

**Figure 4.**
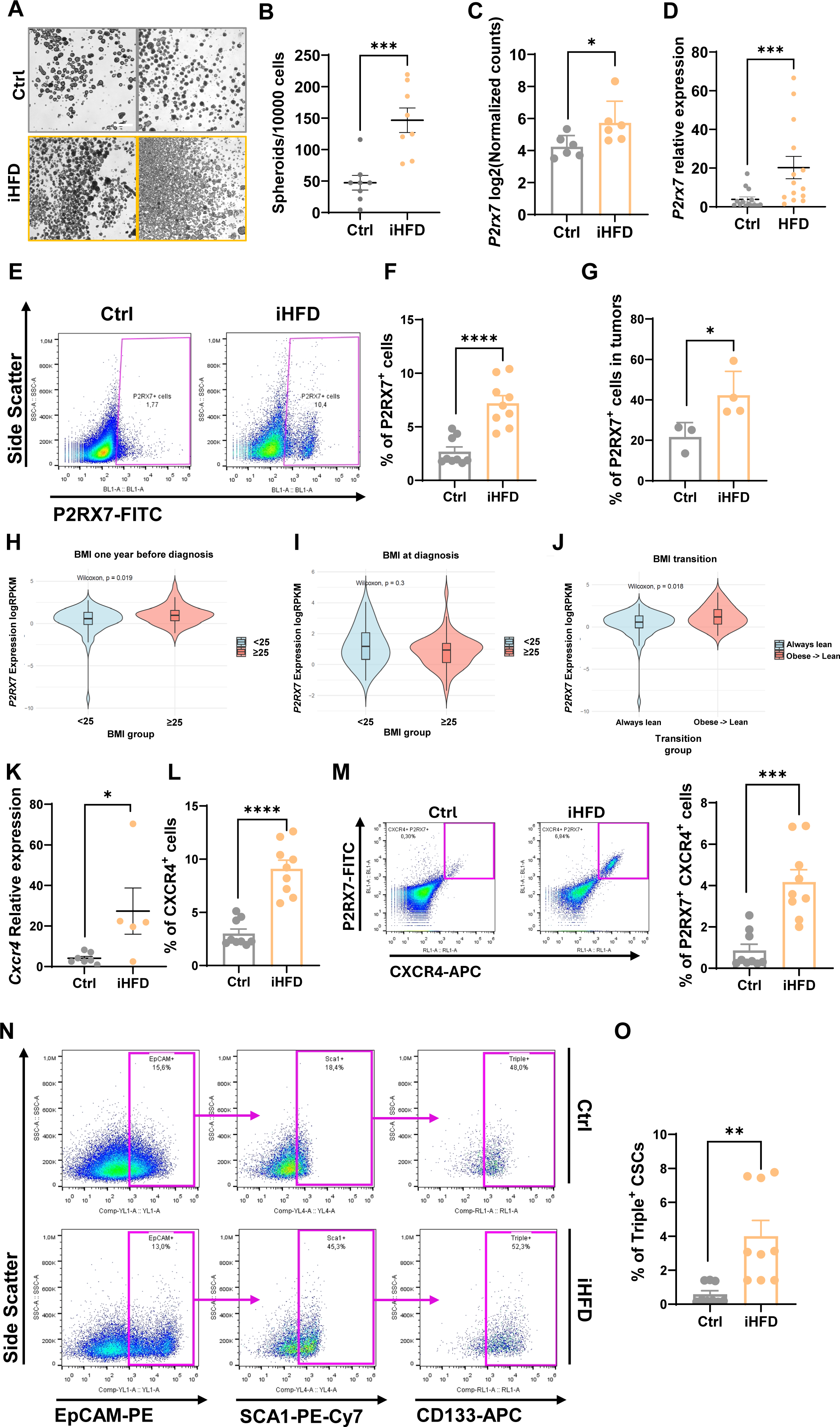
HFD-induced cells present an enrichment in CSC markers and enhanced plasticity. **A)** Representative brightfield microscopy images of spheroids generated from control (Ctrl) and high-fat diet-induced (iHFD) primary tumor cell lines after 5 days in culture. Scale=750 µm. **B)** Quantification of the number of spheroids generated from 10000 cells. Shown is the mean ± SEM (n=8, P values as determined by unpaired T-test). **C)** Quantification of P2rx7 expression in Ctrl and iHFD primary cell lines from the RNA-seq analyses. Shown is the mean ± SEM of log2(normalized counts) (n=6, P values as determined by unpaired T-test). **D)** Relative expression of P2rx7 mRNA levels measured by RT-qPCR in Ctrl and iHFD primary tumor cell lines. Shown is the mean ± SEM (n=14, P values as determined by unpaired T-test). **E-F)** Representative flow cytometry plots showing the levels of P2RX7 surface expression in Ctrl and iHFD cell lines (**E**) and quantification of the expression (**F**), shown as the mean ± SEM (n=3, P values as determined by unpaired T-test). **G)** Quantification of the percentage of P2RX7 positive cells in tumors grown in Ctrl or iHFD-fed mice. Shown is the mean ± STDEV (n=3, P values as determined by unpaired T-test). **H)** Violin plot showing the quantification of P2RX7 expression in PDAC patients based on their BMI 1 year before diagnosis (n for <25=75, n for ≥25=89). P value as determined by Wilcoxon test. **I)** Violin plot showing the quantification of P2RX7 expression in PDAC patients based on their BMI at time of diagnosis (n for <25=35, n for ≥25=54). P value as determined by Wilcoxon test. **J)** Violin plot showing the quantification of P2RX7 expression in PDAC patients based on their BMI transition (patients previously obese that lost their weight versus those that always present a BMI<25) (n for always <25=75, n for obese → lean =35). P value as determined by Wilcoxon test. **K)** Relative expression of Cxcr4 mRNA levels measured by RT-qPCR in Ctrl and iHFD primary tumor cell lines. Shown is the mean ± SEM (n=5, P values as determined by unpaired T-test). **L)** Quantification of CXCR4 expression by flow cytometry in Ctrl and iHFD primary tumor cell lines. Shown is the mean ± SEM (n=3 in triplicates, P values as determined by unpaired T-test). **M)** Representative flow cytometry plots showing the levels of P2RX7/CXCR4 double positive cells in Ctrl and iHFD cell lines (left panel) and quantification of the expression (right panel), shown as the mean ± SEM (n=3, P values as determined by unpaired T-test). **N-O)** Representative flow cytometry plots showing the levels of EpCAM/SCA1/CD133 triple positive cells in Ctrl and iHFD cell lines (**N**) and quantification of the expression (**O**), shown as the mean ± SEM (n=3, P values as determined by unpaired T-test). P values: *, <0.05; **, <0.01; ***, <0.001; ****, <0.0001.

As P2RX7 is considered a CSC-associated marker, we evaluated the presence of other stem-related surface markers in these cell lines. Specifically, we assessed the expression of the “highly metastatic CSC population” marker CXCR4 (36) in iHFD cells and found it to be upregulated at the transcriptomic and protein levels, supporting the link between HFD-induced tumors with a more invasive phenotype (**Figure 4K-L**). Notably, most P2RX7-positive cells in our cultures also co-expressed CXCR4 (**Figure 4M**). We also found an increase abundance of the triple-positive (i.e., EpCAM^+^, SCA1^+^, CD133^+^) stem-like cell population (8) in our iHFD cultures (**Figure 4N-O**). Altogether, these results position HFD as an activator of plasticity and stemness in mouse PDAC. Stem-related markers such as *EPCAM*, *PROM1* or *ISG15* were not differentially expressed between patients with high versus low BMI one year before diagnosis (**Figure S6B-D**). However, *CXCR4* mirrored the expression pattern of *P2RX7* (**Figure S6E-G**), strongly suggesting that HFD may also induce cellular plasticity and a potentially metastatic phenotype in PDAC tumors.

### HFD-induced PDAC in a late adulthood model shows histological and molecular concordance with early adulthood disease

The HFD exposure in an early-adulthood onset revealed an important role of HFD in tumor progression; however, in this model we could not address if the HFD had a role in tumor induction (5, 6). As such, we next investigated whether HFD exposure could play a role in tumor induction in a model resistant to tumor development. To this end, mice were kept with DOX for two months, left for an additional month for DOX clearance before HFD exposure and monitored until one year on a HFD diet or humane endpoint (P60 protocol). This protocol resembles adults that gain mutations due to ageing or other factors and that acquire HFD-like habits later in life (**Figure 5A**). Using this approach, and as expected from our previous work, control mice were resistant to tumor formation (**Figure 5B-E**) (5, 6). Importantly, we found that the HFD intake led to the development of lesions in 63% of the cohort, with 38% of the total mice presenting full PDAC (**Figure 5E**). Next, we performed immunohistopathological analyses of these tumors and found a strong parallelism with the P21 setting, including a highly mesenchymal tumor phenotype characterized by most of the cells expressing vimentin and a very low proportion of cells expressing cytokeratin 19 (**Figure 5F**). iHFD primary tumor-derived cell lines from the P60 protocol shared common features with iHFD P21 cell lines, including a mesenchymal morphology (**Figure 5G**) and a similar expression profile, both at the mRNA and protein levels for P2RX7 and CXCR4 upregulation, as well as a reduction in *Krt19* and *Cdh1* expression (**Figure 5H-I**). Therefore, we found shared common features in the induction and progression of a molecular mechanism in both HFD-exposed P21 and P60 protocols. Furthermore, these results strongly support a critical role for HFD in two important steps of PDAC development, tumor initiation and in tumor progression.

**Figure 5.**
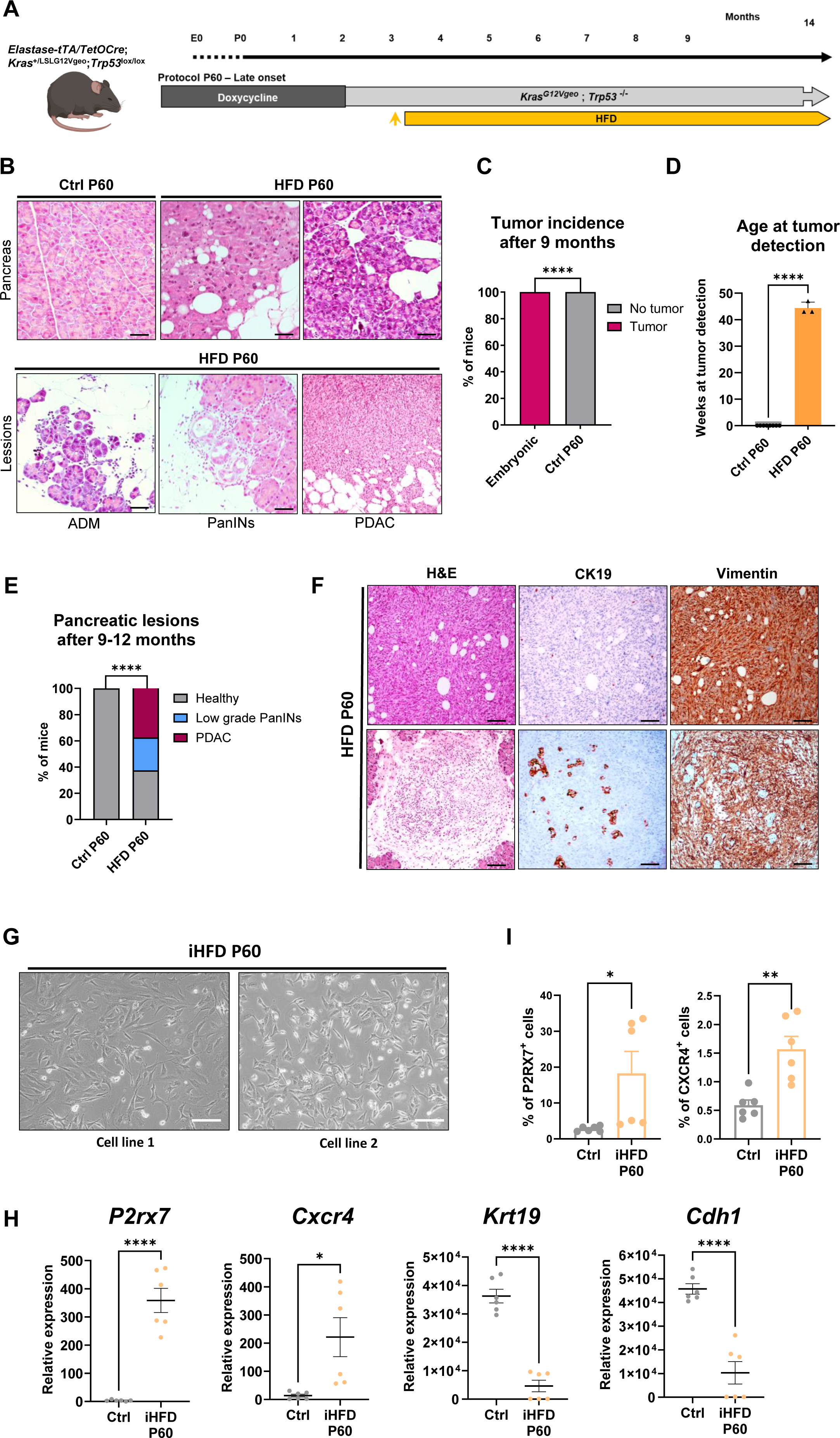
High-fat diet induces tumorigenesis in a late-onset adult PDAC mouse model and presents similar characteristics to the early-onset model. A) Representative scheme of the protocol employed to study the effect of high-fat diet (HFD) in late onset pancreatic cancer. B) Representative brightfield microscopy images of H&E staining from pancreata at different stages of tumor development in the P60 protocol (top panel), as well as the identified pancreatic lesions in the P60 protocol mice fed with HFD. Scale bar= 50 µm. C) Percentage of mice from embryonic (n=19) and P60 controls (Ctrl) (n=8) in which tumors were detected. P values as determined by Chi-square test and Fisheŕs exact test. D) Histogram showing the weeks at tumor detection for the Ctrl (n=8) and HFD-fed (n=3) mice from the P60 protocol. Shown is the mean ± SEM (n=8 mice, P values as determined by One-way ANOVA, Tukey test). E) Percentage of control and HFD P60 mice presenting healthy tissue, low grade PanINs or PDAC (n=8). P values as determined by Chi-square test and Fisher’s exact test. F) Representative brightfield microscopy images of hematoxylin and eosin (H&E) staining, along with cytokeratin 19 (CK19) and vimentin IHC from two tumors generated in P60 HFD mice. Scale bar= 50 µm. G) Representative brightfield microscopy images of primary tumor-derived cell lines from P60 high-fat diet-induced (iHFD) mice. Scale=300µm. H) Relative expression of mRNA levels measured by RT-qPCR of the indicated genes in P21 Ctrl and P60 iHFD primary tumor cell lines. Shown is the mean ± SEM (n=2 in triplicates, P values as determined by unpaired T-test). I) Quantification of P2RX7 (left panel) and CXCR4 (right panel) expression by flow cytometry in P21 Ctrl and iHFD P60 primary tumor cell lines. Shown is the mean ± SEM (n=2 in triplicates, P values as determined by unpaired T-test). P values: *, <0.05; **, <0.01; ***, <0.001; ****, <0.0001.

### HFD-derived tumor cells present enhanced metastatic capacity

Considering the more plastic-like/mesenchymal phenotype of iHFD cell lines, as well as the expression of pro-metastatic-related markers (i.e., CXCR4), we evaluated the capacity of these cells to migrate and invade *in vitro* and *in vivo.* First, we performed wound healing and transwell invasion assays comparing the migratory capacity and invasiveness of control and iHFD cells. In both cases, iHFD cell lines presented an enhanced ability to close wounds and to migrate through Matrigel™ (**Figure 6A-D**). We next evaluated the presence of spontaneous metastases in our GEMMs. As P60 control mice did not form primary tumors or lesions, we focused on the embryonic and P21 protocols. We found that in the P21 protocol, 22.22% of control mice presented liver metastases, while 71.43% of the HFD mice presented both liver and/or lung metastases, showing a pro-metastatic impact of the HFD (**Figure 6E, G**). Due to the aggressive nature and rapid development of primary PDAC tumors in the embryonic model, none of the control mice presented metastases, but in the HFD-exposed embryonic model, we found that 54.54% of mice presented metastases in the liver and lung at humane end point (**Figure 6F-G**). Thus, in general, for both protocols, we found that HFD induced more metastatic burden and primed cells to both the lung and liver (**Figure 6H**).

**Figure 6.**
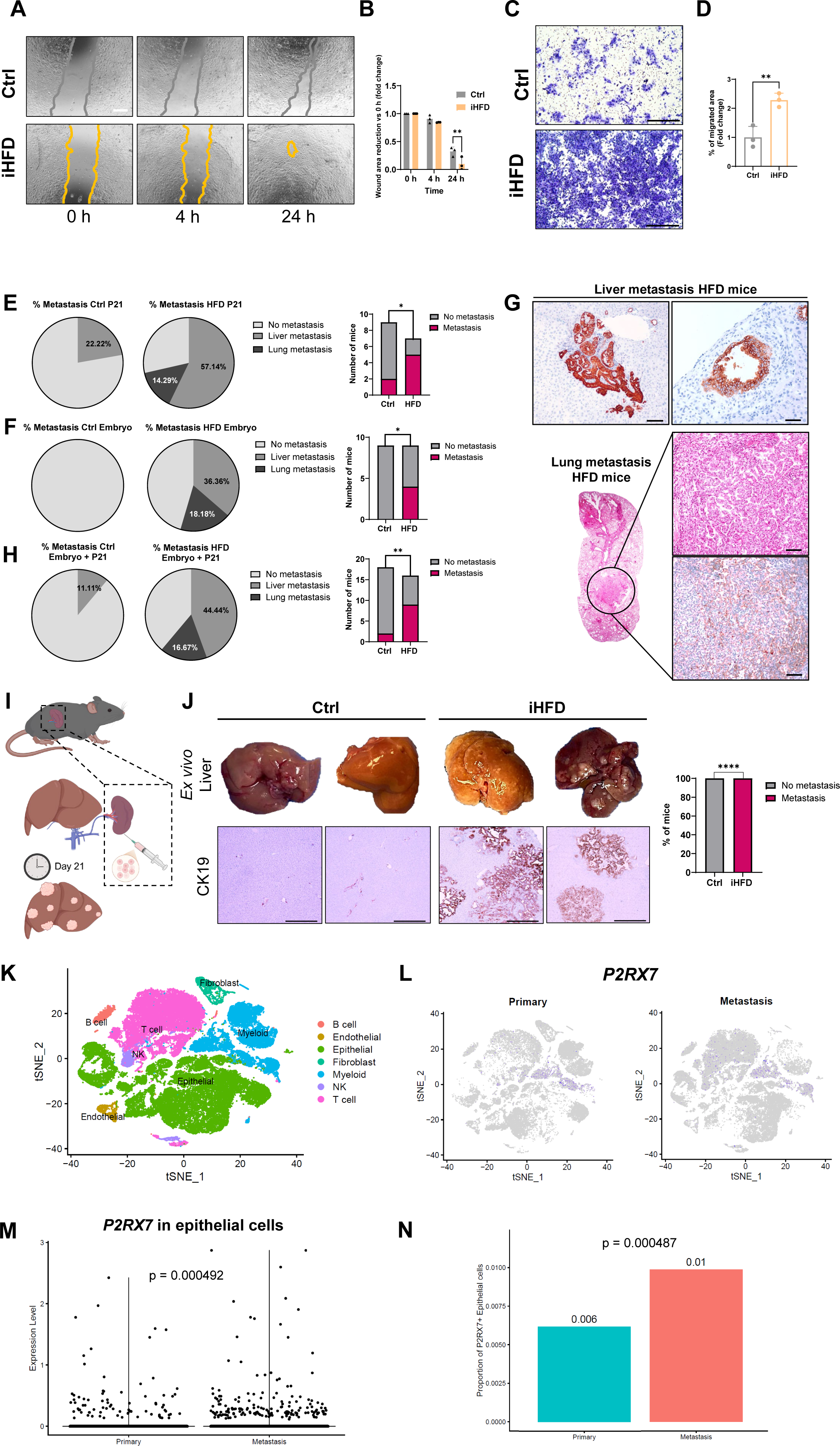
High-fat diet induces a pro-metastatic phenotype. **A)** Representative brightfield microscopy images of wound healing assays at 0-, 4-and 24-hours post-scratch in both control (Ctrl) and high-fat diet-induced (iHFD) cell lines. Scale=750 µm. **B)** Quantification of wound area reduction shown in A represented as the mean fold change ± STDEV with time 0h set as 1.0 (n=3, P values as determined by unpaired T-test). **C)** Representative brightfield microscopy images of crystal violet-stained transwell migration assays at 24 hours post-seeding in both Ctrl and iHFD cell lines. **D)** Quantification of migration area shown as the mean fold change ± STDEV with Ctrl cell lines set as 1.0 (n=3, P values as determined by unpaired T-test). **E)** Left panel: Proportion plots showing the percentage of Ctrl and iHFD mice with metastasis from the P21 protocol. Right panel: Quantification of the number of mice with metastasis in both conditions (n=9 in Ctrl and 7 in HFD, P values as determined by Chi-square test). **F)** Left panel: Proportion plots showing the percentage of Ctrl and HFD mice with metastasis from the embryonic (Embryo) model. Right panel: Quantification of the number of mice presenting metastasis in both conditions (n=9 in Ctrl and 9 in HFD, P values as determined by Chi-square test). **G)** Representative brightfield microscopy images from cytokeratin 19 immunohistochemistry-stained and H&E-stained samples from metastatic disease in HFD mice from the P21 protocol in liver (top panel) and lung (bottom panel). Scale bar= 500 µm. **H)** Left panel: Proportion plots showing the percentage of Ctrl and HFD mice from both models (embryonic and adult P21), which present metastasis. Right panel: Quantification of the number of mice presenting metastasis in both conditions (n=18 in Ctrl and 16 in HFD, P values as determined by Chi-square and Fisheŕs exact test). **I)** Graphical scheme showing the basis of the intrasplenic assay to evaluate the liver metastatic capacity of a cell line. **J)** Left panel: Representative pictures of livers from the intrasplenic assay injected with Ctrl and iHFD cell lines (top) and representative brightfield microscopy images from cytokeratin 19 (CK19) immunohistochemistry of Ctrl and HFD livers (bottom). Right panel: Quantification of the percentage of mice that showed macroscopic metastasis in the intrasplenic assay (n=3 cell lines per condition, P values as determined by Chi-square and Fisheŕs exact test). **K)** tSNE plot showing the different annotated cell types from the Liu et al. (38) single cell RNA-seq dataset of human primary tumors and metastasis. **L)** tSNE plots showing the P2RX7-expressing (blue) cells in the Liu et al. dataset separated by condition (primary for primary tumor and metastasis for liver metastasis). **M)** Quantification of P2RX7 expression in the epithelial compartment in both primary tumor and metastasis. P value indicated and determined by Wilcoxon test. **N)** Bar plot of P2RX7-expressing cells in primary tumors and metastasis. P value indicated and determined by Fisher test. P values: *, <0.05; **, <0.01; ***, <0.001; ****, <0.0001.

To further validate that the pro-metastatic effect of HFD is imprinted in tumor cells, we performed an intrasplenic experimental metastasis assay in C57Bl/6 mice with control and iHFD cell lines to evaluate their capacity to generate liver metastasis (**Figure 6I**). After three weeks, we sacrificed the mice and found that all iHFD cell lines colonized the liver and produced metastasis, while the controls cell lines did not (**Figure 6J**). Considering that HFD-induced fatty liver could be a better soil for metastatic seeding (37), we evaluated the capacity of control cell lines to metastasize in mice fed with normal diet or HFD (**Figure S7A-B**). Although we did not find an increased metastatic burden in HFD-fed mice (**Figure S7C-D**), we observed that the EpCAM^+^ cells found in the liver were mainly P2RX7^+^, suggesting the implication of this receptor in metastatic seeding (**Figure S7E**). To validate this observation in a human setting, we interrogated an already annotated single cell RNA-seq dataset comprising 12 matched samples from primary tumors and liver metastases (38) (**Figure 6K**). *P2RX7* expression was mainly restricted to the immune compartment both in primary tumor and metastasis (**Figure 6L**); however, we found a small proportion of cells in the epithelial compartment expressing the gene, as expected from for CSC marker (**Figure 6M**). In line with our results in mouse models, we found that metastasis presented a higher proportion of P2RX7-expressing tumor cells compared to primary tumors (**Figure 6N**). Altogether, these results support that HFD induces a metastatic phenotype in tumor cells characterized by P2RX7-expression during tumor evolution.

### Lipid-conditioned macrophages drive plasticity in tumor cells through a CAMP-P2RX7 axis

We next studied the mechanisms by which HFD induces the mesenchymal/plastic phenotype of tumor cells. Since fatty acids (FAs) have been reported to induce a pro-CSC phenotype in human tumor cells (39), we asked whether increase exposure to FAs could cause the observed phenotype. Regarding P2RX7 expression, none of the FAs we tested (oleic acid, monounsaturated; linoleic acid, polyunsaturated; palmitic acid, saturated) could increase P2RX7 levels in mouse control cell lines (**Figure 7A**) or in PDX-derived human primary cell lines (**Figure 7B**). Thus, we hypothesized that the increase in P2RX7 and the phenotype induction could be driven by components of the tumor microenvironment such as adipose tissue or neutrophils (9). To test this hypothesis, we cultured adipose tissue from regular diet and HFD-fed C57Bl/6 mice and obtained adipose tissue conditioned medium (ACM). Exposure of control cell lines to ACM did not result in an increase in P2RX7 (**Figure 7C**). Also, we found that neutrophils were poorly infiltrated or almost absent in HFD tumors (**Figure S8A-B**). Next, we assessed highly tumor infiltrating macrophages, although they were not significantly different in both settings (**Figure S8A, C**). Importantly, since the human RNA-seq data revealed an enrichment in gene signatures for lipid-conditioned macrophages and antimicrobial response, we theorized that macrophages might influence *P2RX7* expression in tumor cells and activate it via its ligand, CAMP. Indeed, *P2RX7* and *CAMP* positively correlated in human PDAC (**Figure 7D**) and CAMP expression has been linked to metabolic diseases related to obesity such as atherosclerosis or diabetes (40, 41). To address this, we cultured immortalized primary bone marrow-derived macrophages (IBMDMs) with FAs, and measured *Camp* expression at the transcriptomic level. We found that naive IBMDMs did not express *Camp*, whereas FAs stimulation robustly induced its expression (**Figure 7E**). Importantly, mice fed with HFD presented increase CAMP-expressing macrophages in the bone marrow (BM) and intratumorally compared to control mice (**Figure 7F-G**). As it was described that macrophage-derived hCAP-18/LL-37 can induce P2RX7 expression in human cell lines (26), we next cultured control cells with naive macrophages conditioned medium (MCM) or FAs-treated macrophage conditioned medium (F-MCM). Interestingly, F-MCM induced more the expression levels of *P2rx7* compared to control cell lines treated with MCM (**Figure 7H**). Furthermore, F-MCM decreased *Epcam* expression levels (**Figure 7I**) and induced a more mesenchymal phenotype (**Figure 7J**), supporting that FAs derived from the diet conditioned macrophages to produce molecules, such as CAMP, that induce P2RX7 expression and changes in the cellular phenotype. Notably, stimulation of control cell lines with recombinant CAMP (rCAMP) increased the levels of P2RX7 and CXCR4 (**Figure 7K-L**). In addition, we evaluated whether rCAMP could also account for the elevated expression of PGLYRP1 observed in iHFD cells (**Figure S9A**). We found that rCAMP significantly increased PGLYRP1 expression levels in control cell lines (**Figure S9B**). Due to the protective role of PGLYRP1 blocking macrophage phagocytosis in PDAC cells (8), we next performed a phagocytosis assay to determine whether iHFD cells exhibit enhanced protection and immune evasive features. Consistent with the overexpression of PGLYRP1, iHFD cell lines were phagocytosed less efficiently by primary macrophages than control cell lines (**Figure S9C-D**).

**Figure 7.**
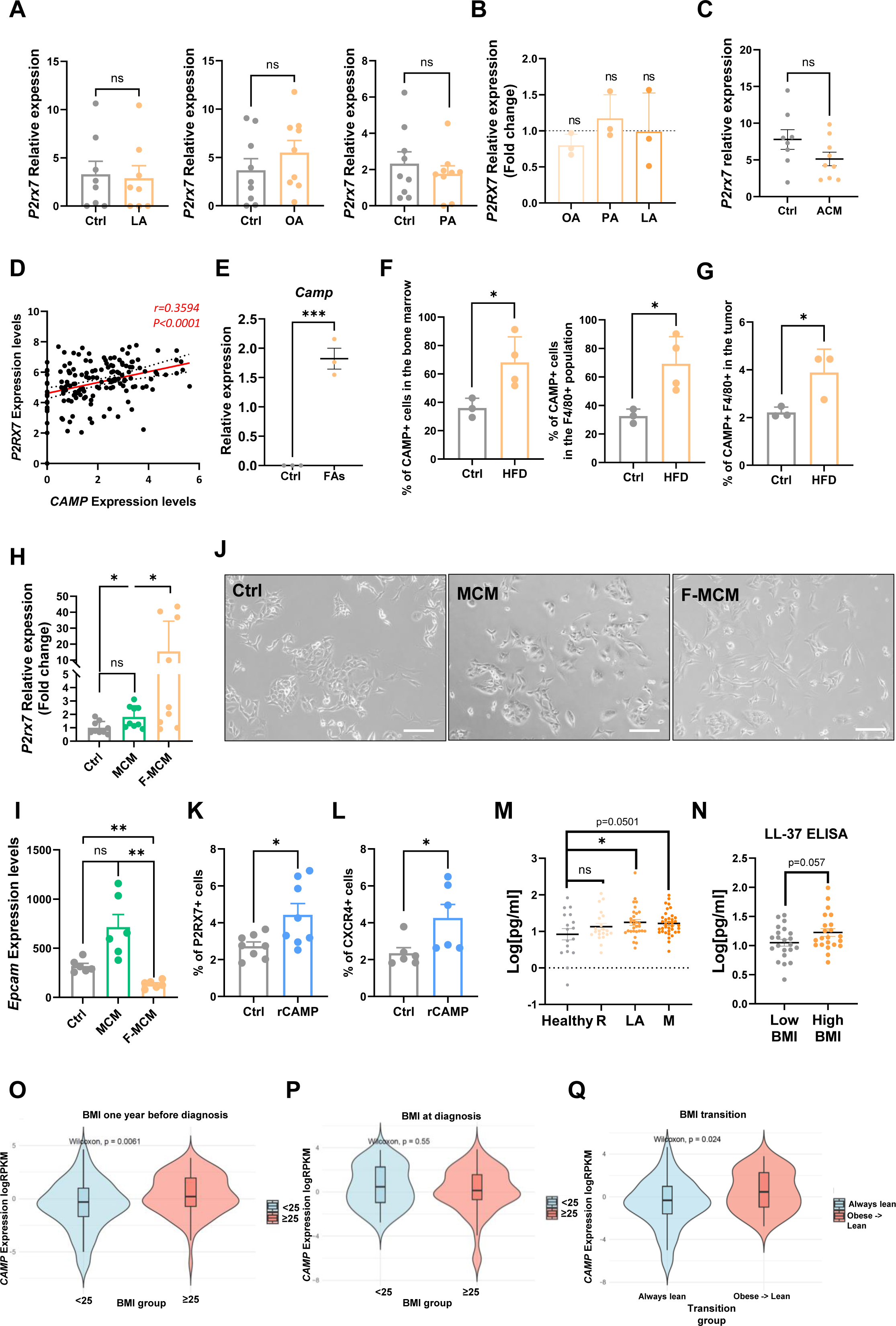
P2RX7-CAMP axis is induced by macrophages. **A)** Quantification of P2rx7 expression by RT-qPCR in control (Ctrl) murine cell lines treated with fatty acids (FAs) (linolenic acid, LA [200µM]; oleic acid, OA [50µM]; palmitic acid, PA [50µM sodium palmitate]). Shown is the mean ± SEM (n=4 cell lines in duplicates for LA, and 3 cell lines in triplicate for OA and PA, P values as determined by unpaired T-test). **B)** Quantification of P2RX7 expression by RT-qPCR in PDX-derived primary human cell lines treated with FAs. Shown is the mean fold change ± SEM with controls set as 1.0 (n=3 cell lines, P values as determined by unpaired T-test). **C)** Quantification of P2rx7 expression by RT-qPCR in Ctrl murine cell lines treated with adipose tissue conditioned medium (ACM) for 48 hours. Shown is the mean ± SEM (n=4 cell lines in duplicates, P values as determined by unpaired T-test). **D)** Dot plot showing the correlation between P2RX7 and CAMP expression in TCGA patients. P values and r were calculated by Pearson’s correlation. **E)** Quantification of Camp expression by RT-qPCR in immortalized bone marrow-derived macrophages treated with FAs (linolenic acid, LA; oleic acid, OA; palmitic acid, PA). Shown is the mean ± STDEV (n=3, P values as determined by unpaired T-test). **F)** Quantification of CAMP positive total cells (left panel) and CAMP positive cells in macrophages (F4/80) (right panel) from the bone marrow determined by flow cytometry in mice fed with Ctrl diet and HFD. Shown is the mean ± STDEV (n=3 mice for control group, 4 for HFD group, P values as determined by unpaired T-test). **G)** Quantification of CAMP positive tumor-associated macrophages determined by flow cytometry in mice fed with Ctrl diet and HFD. Shown is the mean ± STDEV (n=3 mice, P values as determined by unpaired T-test). **H)** Quantification of P2rx7 expression by RT-qPCR in Ctrl murine cell lines treated with macrophage conditioned medium (MCM) or fatty acid-trained macrophage conditioned medium (F-MCM) for 48 hours. Shown is the mean fold change ± SEM with Ctrl set as 1.0 (n=3 cell lines in triplicates, P values as determined by One Way ANOVA). **I)** Quantification of Epcam expression by RT-qPCR in Ctrl murine cell lines treated with MCM or F-MCM for 48 hours. Shown is the mean fold change ± SEM with Ctrl set as 1.0 (n=3 cell lines in duplicates, P values as determined by One Way ANOVA). **J)** Representative brightfield microscopy images of Ctrl cells treated with MCM and F-MCM. Scale bar= 50µm. **K)** Quantification of P2RX7 positive cells determined by flow cytometry in a Ctrl cell culture treated with recombinant (r) CAMP (100 µg/mL) for 48 hours. Shown is the mean ± SEM (n=8, P values as determined by unpaired T-test). **L)** Quantification of CXCR4 positive cells determined by flow cytometry in a Ctrl cell culture treated with rCAMP (100 µg/mL) for 48 hours. Shown is the mean ± SEM (n=6, P values as determined by unpaired T-test). **M)** Quantification by ELISA of soluble LL-37 levels in sera from healthy donors (Healthy) (n=17) and PDAC patients with resectable (R) (n=21), locally advanced (LA) (n=30) or metastatic disease (M) (n=37). Shown is the mean of log normalized concentration ± SEM (P values as determined by One-Way ANOVA, Dunnet’s test). **N)** Quantification by ELISA of soluble LL-37 levels in sera from PDAC patients from the highest and lowest quartiles of BMI (n=22 patients per condition). Shown is the mean of log normalized concentration ± SEM (P values as determined by unpaired T-test). **O)** Violin plot showing the quantification of CAMP expression in PDAC patients based on their BMI 1 year before diagnosis (n for <25=75, n for ≥25=89). P value as determined by Wilcoxon test. **P)** Violin plot showing the quantification of CAMP expression in PDAC patients based on their BMI at time of diagnosis (n for <25=35, n for ≥25=54). P value as determined by Wilcoxon test. **Q)** Violin plot showing the quantification of CAMP expression in PDAC patients based on their BMI transition (patients previously obese that lost their weight versus those that always present a BMI<25) (n for always <25=75, n for obese → lean =35). P value as determined by Wilcoxon test. P values: *, <0.05; **, <0.01; ***, <0.001; ****, <0.0001.

After finding an increase in CAMP expression in murine BM macrophages, we evaluated its expression in patients’ PBMCs as a possible proxy marker for HFD-induced PDAC. Although macrophages are not present in circulation, we detected *CAMP* expression in PBMCs from both patients and non PDAC controls (**Figure S10A-B**). As this circulating CAMP is unlikely to be macrophage derived, no significant differences were observed between groups stratified by BMI. However, in both groups (BMI\25 and BMI^25), PDAC patients exhibited higher *CAMP* expression in PBMCs compared with non-PDAC controls (**Figure S10C-D**). As a complementary approach, we assessed the serum levels of the hCAP-18-derived peptide LL-37 in a cohort of healthy donors and PDAC patients. We found that PDAC patients with locally advanced and metastatic disease presented increased levels of sera LL-37, compared with healthy donors (**Figure 7M**). When we divided PDAC patients according to their BMI and compared the highest and lowest quartiles of the cohort, we found higher levels in those with high BMI (**Figure 7N**). These distinctions became more evident upon assessing *CAMP* expression at the transcriptomic level in patients, where we observed a parallel trend with *P2RX7* and *CXCR4* expression; specifically, increased levels among individuals with high BMI one year prior to diagnosis (**Figure 7O-Q**). Finally, we also evaluated EpCAM and hCAP-18/LL-37 levels by immunohistochemistry in a small patient cohort divided by BMI. In line with our results, patients with BMI>25 displayed lower EpCAM expression in tumor cells and higher levels of LL-37 within the immune compartment compared to patients with BMI<25 (**Figure S11A-C**). Altogether, these results support that HFD-induced obesity induces the systemic expression of CAMP by macrophages, which may contribute to the induction of an immune evasive and more mesenchymal plastic tumor cell state within the tumor.

## Discussion

Mouse models of PDAC have been instrumental over the past decade for preclinical studies, enabling the field to capture the complexity of a disease in which the tumor microenvironment plays a central role. Among them, the KPC model (*Pdx1/Ptf1a*-Cre; *Kras^+/^*^G12D^; *Trp53*^mut^ ^or^ ^KO^) has been widely used (42), yet, like all models, it presents inherent limitations. The acquisition of oncogenic mutations during embryonic development generates tumors that mimic key features of human PDAC. However, in humans, these mutations are not acquired during embryonic development and this could be the reason why it has been suggested that mutations are not sufficient to drive PDAC (43–45). Indeed, Carpenter *et al.* recently demonstrated that healthy individuals frequently harbor *KRAS*-mutant PanINs (4), and the prevalence of these lesions far exceeds the incidence of PDAC. This strongly suggests that oncogenic potential does not reside solely in mutations but requires additional events, particularly chronic inflammation, to initiate tumorigenesis.

Multiple studies over the past two decades have linked inflammatory cues to PDAC initiation and progression, whether through epigenetic priming (46, 47), microenvironmental remodeling (48, 49) or, as our group previously showed, through tissue damage that propagates oncogenic signaling and suppresses senescence (5, 6). To dissect these processes, we developed inducible mouse models in which mutations arise specifically in the acinar compartment of adult mice using a Tet-Off system. This KPeC model (*Elas-tTA/tetO-Cre*; *Kras^+/LSL^*^G12Vgeo^; *Trp53*^lox/lox^) revealed that the adult pancreas is intrinsically resistant to oncogenic transformation and that chronic pancreatitis is required to overcome this barrier, a concept now widely accepted in the human setting. Importantly, this model provides a unique opportunity to interrogate “early-onset” and “late-onset” PDAC, which we modelled through our P21 and P60 protocols.

The rising incidence of PDAC in individuals under 50 years of age (3) suggests that environmental exposures, including diet, may play a major role. By inducing mutations at different ages, we observed that younger hosts were more permissive to tumor formation than the adult pancreas, which was completely resistant to transformation by the *Kras* oncogene expression and *Trp53* loss, as previously reported (5). This may reflect the retention of plasticity or stem-like features within subsets of acinar cells, traits that become progressively restricted with age (50–52).

Strikingly, we found that HFD consumption increased tumor aggressiveness, metastatic behavior, and reduced survival in the P21 protocol. These findings led us to hypothesize that chronic HFD exposure, particularly during childhood or early adulthood, creates a systemic inflammatory landscape that facilitates oncogenic transformation once mutations arise. Supporting this idea, we observed that HFD-induced systemic signaling imprints stable transcriptional changes in tumor cells that persist in culture, likely through epigenetic mechanisms. The acquisition of a plastic, stem-like phenotype, robustly preserved in iHFD cell lines, has important clinical implications, as such states are strongly associated with therapy resistance and metastatic behavior (53–56). Notably, this phenotype has not been previously described in HFD-fed PDAC models.

Noteworthy, in our late-onset protocol (P60) we could induce preneoplastic lesions and full tumors in a model that is normally resistant to the disease. This shows for the first time a link between specific dietary habits and tumor induction in a model that presents oncogenic mutations at this age stage. In alignment with our results with the P21 protocol, HFD-induced tumors and tumor cell lines both presented a mesenchymal histology and morphology, as well as an enrichment in plastic-like cells characterized by the expression of P2RX7 and CXCR4. These results strongly support the effect of diet in the molecular characteristics and dynamics of PDAC formation and evolution despite the age of mutation acquisition.

Although other inducible KRAS models have been used to study the intersection between HFD and PDAC, they often rely on protocols that do not account for age-dependent differences in acinar cell susceptibility to transformation (9, 10). These studies attributed the tumor-promoting effects of HFD to enhanced desmoplasia, collagen deposition, and inflammation mediated by COX2 and IL-1β, implicating pancreatic stellate cells, neutrophils, and adipocytes as key drivers. In contrast, our models did not support a major role for neutrophils or adipose tissue. Instead, macrophages emerged as central regulators of the tumor cell state observed both *in vitro* and *in vivo*. These discrepancies may reflect differences in model design or the extended tracking period in our study, which spanned up to one year. Importantly, the morphological and transcriptional reprogramming toward a mesenchymal/plastic-like state had not been previously reported. Moreover, our analysis of real-world patient data corroborated these findings, reinforcing their clinical relevance. Collectively, these observations highlight HFD-induced obesity as a critical variable that should be considered when studying interpatient PDAC heterogeneity and raises the question of whether dietary interventions could reverse or mitigate this acquired phenotype.

Mechanistically, we show that FAs systemically condition macrophages to produce CAMP. In humans, macrophage-derived hCAP-18 (the human homolog of murine CAMP) and its active peptide LL-37 have been shown to activate the CSC compartment in PDAC and contribute to tumorigenesis (26). We found that both CAMP and its receptor P2RX7 were upregulated in patients with high BMI and in HFD-derived tumors. Furthermore, CAMP induced the expression of PGLYRP1, another antimicrobial protein produced by neutrophils and CSCs that protects tumor cells from macrophage-and T-cell-mediated elimination (8). This CAMP-P2RX7-PGLYRP1 axis not only reinforces the plastic, stem-like state of HFD-induced tumor cells but also shields them from immune clearance and promotes metastatic behavior.

Consistent with this, we found that HFD enhances metastasis in a tumor cell-intrinsic manner. In this context, the “seed” acquires the capacity to colonize distinct environments (lung and liver) independently of the host, as HFD-fed mice did not exhibit increased metastasis when injected with control cell lines in an intrasplenic assay. Notably, metastatic EpCAM⁺ tumor cells consistently expressed high levels of P2RX7, underscoring its importance for metastatic success, a role previously described in melanoma and prostate cancer (57, 58). Interestingly, in colorectal cancer the P2RX7-CAMP axis appears to function in the opposite direction (59), highlighting the context-specific nature of this pathway.

Although further research is needed, our findings position P2RX7 as a central driver of HFD-and obesity-related PDAC, opening the door to more personalized therapeutic strategies for this patient subgroup. Circulating hCAP-18/LL-37 may be a potential biomarker for early PDAC detection and identifying candidates for targeted therapy. However, our analysis showed minor differences in LL-37 serum levels compared to RNA-seq results. This phenomenon may be attributed to the possibility that patients with elevated BMI one year prior to diagnosis are misclassified as having low BMI at the time of diagnosis due to disease-related weight loss. Thus, proper clinical data management seems to be crucial to properly assess these changes.

In conclusion, here we describe the effects of HFD in tumor development in novel early onset and late onset protocols for PDAC GEMMs. Our results show that diet could influence the transcriptional state of tumor cells, rendering them a more stem/plastic-like state, which favors the metastatic disease. HFD induces the expression of P2RX7, which is activated via macrophage-derived CAMP and favors this characteristic tumor cell state. In patients, high BMI correlates with *P2RX7* and *CAMP* expression, as well as the metastatic marker *CXCR4*, and it is preserved despite weight loss during disease evolution. Altogether, these findings not only reveal a diet-and obesity-driven mechanism that shapes tumor cell plasticity and metastatic potential but also highlights P2RX7 and CAMP as actionable biomarkers and therapeutic targets, paving the way for early detection, risk stratification, and personalized interventions in PDAC patients.

### Lead contact

Requests for further information and resources of this study should be directed to the lead contact, Dr. Carmen Guerra.

## ACKNOWLEDGEMENTS

We thank the patients and healthy controls for donating samples. We especially thank the help of Marta San Roman, María del Carmen Lechuga, Alejandra López, and Raquel Villar from the CNIO Experimental Oncology group. We also want to thank the CNIO Animal Facility, IIBm Microscopy Unit, CNIO Confocal Microscopy Unit, CNIO and IJC Genomics Unit and the CNIO Histopathology Unit for their support. A.G.d-R. was supported by an AECC Predoctoral Fellowship (PRDMA246123GALV) and J.C.L-G. was supported by a “CNIO Friends” Postdoctoral Fellowship and an AECC Postdoctoral Fellowship (POSTD258037LÓPE). E.Z.-D. and P.S. were supported by “Formación de Personal Investigador” (FPI) fellowship (PRE2022-102952) from the Spanish Ministry of Sciences and Innovation and by a fellowship from the China Scholarship Council respectively. This work has been supported by the IX Carmen Delgado/Miguel Pérez-Mateo grant (C.G.), a CRIS Cancer Foundation grant, European Research Council (ERC-AG/695566-THERACAN), CIBERONC Fund (CB21/12/00121), and the Agencia Estatal de Investigación cofounded with the European Regional Development Fund, ‘A way to make Europe’ project PID2024-160242NB-I00 (C.G., M.B.). Also, this work has received support from the Cancer Research Institute (New York, NY, USA, B.S.,Jr.), the AECC with a LAB AECC grant (LABAE223389SANC, to P.S.), the AIRC (IG 26201, G.C.; AIRC IG 32351, C.C.; MBe is supported by AIRC fellowships, 28054, 29829), the PNRR Program (code PNRR-MCNT2-2023-12377229, CUP C53C23001140007) and NEXTGENERATION-EU Project “ICSC” SPOKE8 (CN00000013, CUP J33C22001180001) (both C.C.; PRIN 2022, code: 2022CMMRWA, CUP: B53D23008010006, C.L.), Fondazione Italiana Malattie Pancreas (FIMP-Ministero Salute J38D19000690001, C.L.), and the European Regional Development Fund, ‘A way to make Europe’ ERDF project PID2024-159192OB-I00 (M.E.) and PID2024-157515OB-I00 (B.S.,Jr.). S.P.P. and P.A. are funded by Cancer Research UK (grant C12077/A26223) and Pancreatic Cancer UK, with patient serum samples provided through the PCUK ADEPTS study (REC reference: 06/Q0512/106). S.P.P. is also supported by the National Institute for Health and Care Research University College London Hospitals Biomedical Research Centre. We also want to thank Dr. Mercedes Rodriguez Garrote, Marta Pardo, and Biobank and Biomodels Platform (PT20/0045; PT23/00098), ISCIII research and development platforms in biomedicine and health sciences for supplying the clinical samples and data, as well as the UCL Genomics (UCLG), Zayed Centre for Research into Rare Disease in Children, London, UK (Research Resource Unique Identifier: RRID:SCR_027010) for help with the PBMC RNA-seq dataset. This work has also been developed upon work from COST Action “Identification of biological markers for prevention and translational medicine in pancreatic cancer (TRANSPAN),” CA21116, supported by COST (European Cooperation in Science and Technology).

Finally, we would like to thank the other current and past members of the CNIO Experimental Oncology group and the Cancer Stem Cells and Fibroinflammatory Microenvironment group for helpful discussions, technical assistance and support, as well as María del Castillo and Marta Hergueta from the CNIO Microenvironment and Metastasis group, and Jaime Martínez from CNIO Epithelial Carcinogenesis group.

## AUTHOR CONTRIBUTIONS

Conceptualization of the study: C.G. and M.B. *In vitro* experimental work: A.G.-d-R., J.C.L.-G., S.A., J.E., B.P.-A., A.R., P.Sun. *In vivo* experimental work: A.G.-d-R., J.C.L.-G., P.Sun., E.Z.-D., B.R.-P., R.B., S.J.-P. Bioinformatic analyses: A.G.-d-R., J.C.L.-G., A.A., A.N., K.F. Mouse pathology analyses: A.G.-d-R., J.C.L.-G., C.G. Human pathology analyses: M.Be., C.L. Human transcriptomic data from patients: L.A., G.C. Formal analyses and investigation were made by A.G.-d-R., J.C.L.-G. Resources were provided by B.S.Jr., C.G., M.B. The original draft was written by A.G.-d-R. and J.C.L.-G. All authors contributed reviewing and editing the manuscript. C.G. and M.B supervised the development of the project. P.S., G.C., M.E., S.P.P., P.A., C.C., B.S.Jr., C.G. and M.B. obtained the funding involved in the development of the study.

## Notes

### Competing Interest Statement

The authors have declared no competing interest.

### Summary of Updates

After conversation with a group of collaborators, we have edited the title and made discrete changes in the text. We also provide new supplementary material.

