## Supplementary figures and tables for "High-fat diet imprints a macrophage-tumor CAMP-P2RX7 axis driving pancreatic cancer initiation, plasticity and metastasis"

#### *Supplementary data*

### SUPPLEMENTARY FIGURES

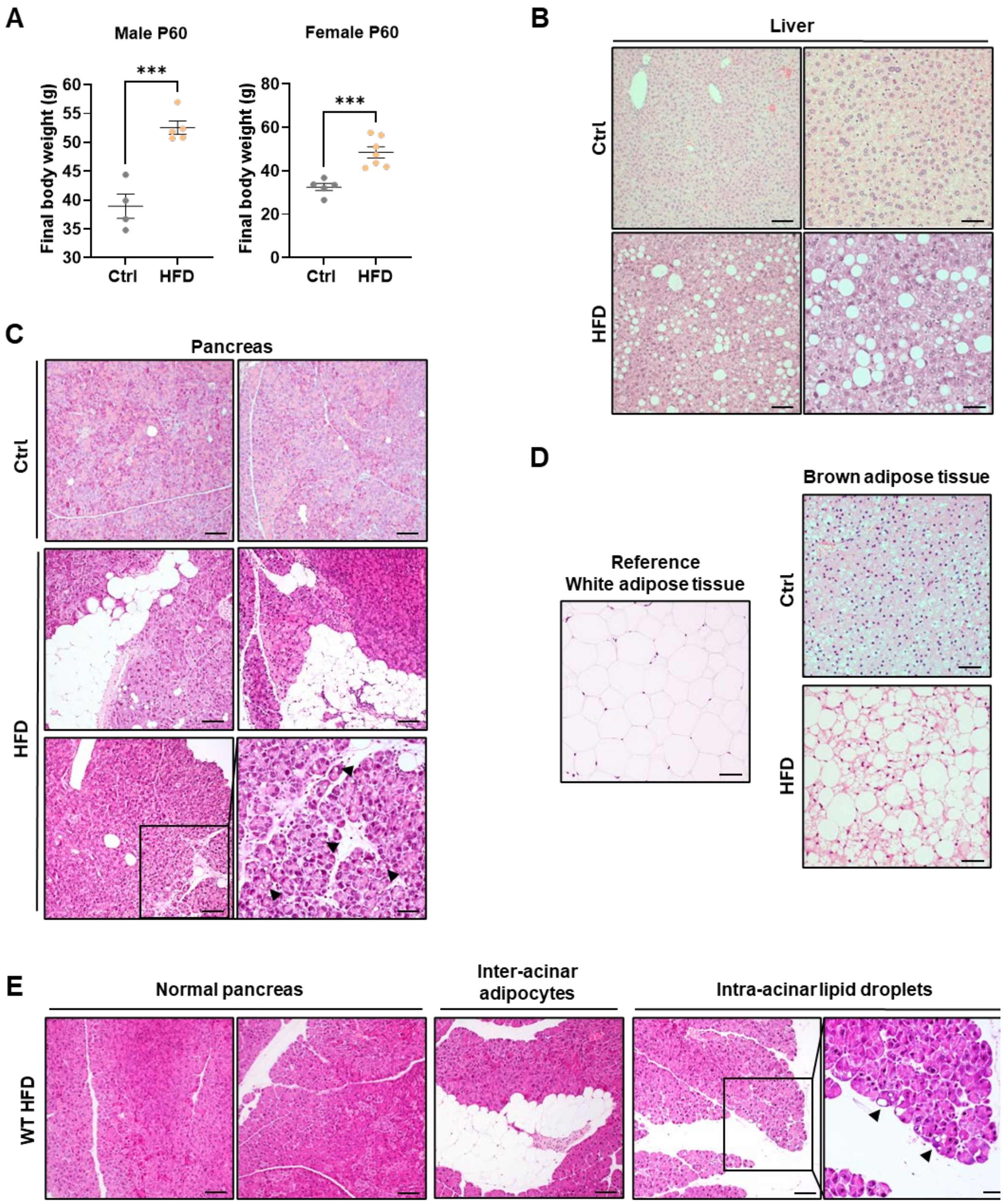


***Figure S1. High-fat diet induces obesity traits in murine models. A)*** *Quantification of final weight from control (Ctrl) and high-fat diet (HFD) exposed mice included in the P60 protocol divided by sex (males, left panel; females, right panel), shown as the mean ± SEM (n=4 Ctrl males, 5 HFD males, 5 Ctrl females, 7 HFD females, P values as determined by unpaired T-test).* ***B)*** *Representative brightfield microscopy images of H&E staining from the liver from Ctrl and HFD exposed mice.* ***C)*** *Representative brightfield microscopy images of H&E staining from the pancreata from Ctrl and HFD exposed mice. Scale bar=100 µm (left column), 50 µm (right column). Black arrowheads point out intra-acinar lipid droplets observed in HFD-fed mice.* ***D)*** *Representative brightfield microscopy images of H&E staining from the white (as a reference) and brown*

*adipose tissue from Ctrl and HFD exposed mice.* ***E)*** *Representative brightfield microscopy images of H&E staining from the pancreata from C57Bl/6 mice exposed to HFD. Black arrowheads point out intra-acinar lipid droplets. Scale bar=100 µm, 50 µm (picture in the right).*

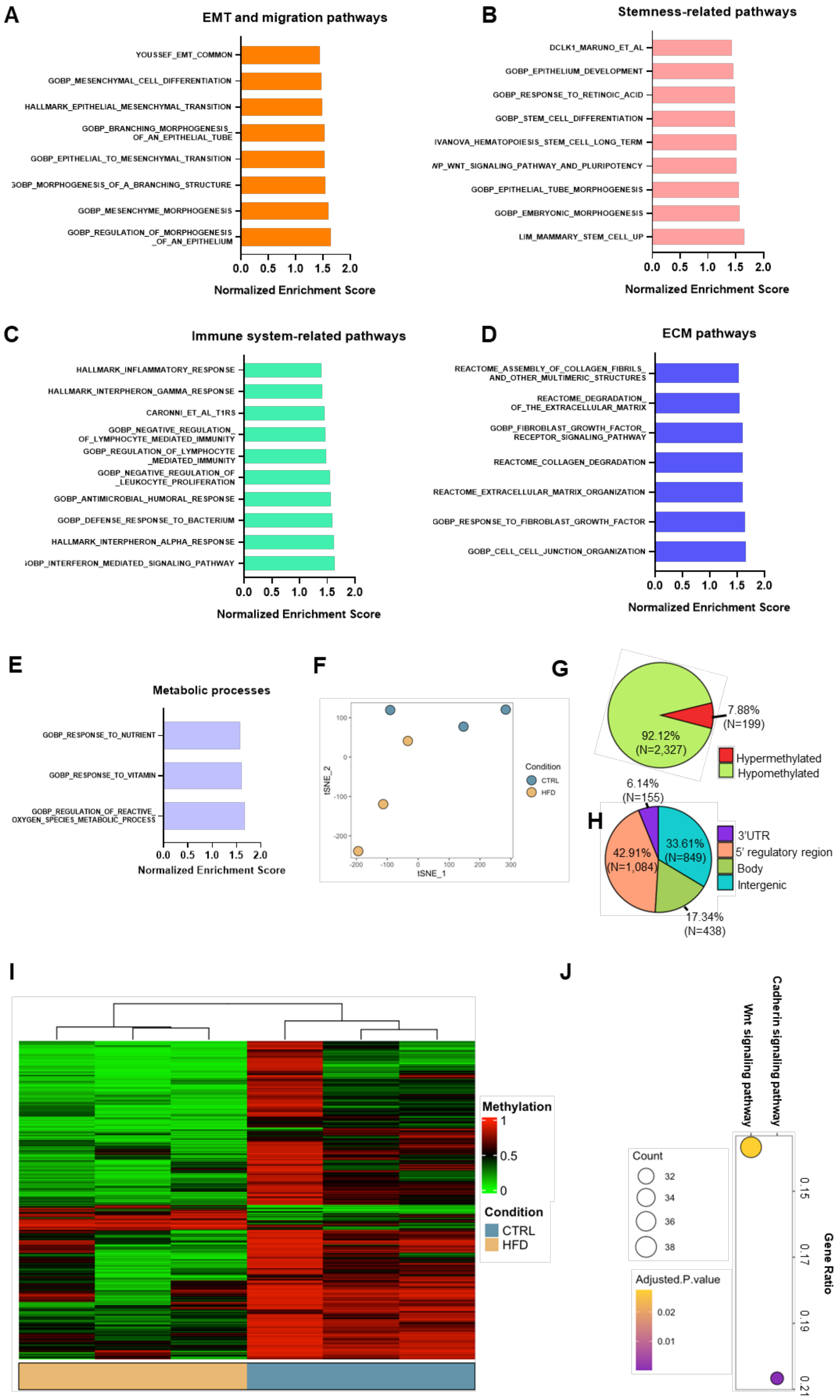
A
EMT and migration pathways
YOUSSEF_EMT_COMMON
GOBP_MESENCHYMAL_CELL_DIFFERENTIATION
HALLMARK_EPITHELIAL_MESENCHYMAL_TRANSITION
GOBP_BRANCHING_MORPHOGENESIS_OF_AN_EPITHELIAL_TUBE
GOBP_EPITHELIAL_TO_MESENCHYMAL_TRANSITION
GOBP_MORPHOGENESIS_OF_A_BRANCHING_STRUCTURE
GOBP_MESENCHYME_MORPHOGENESIS
GOBP_REGULATION_OF_MORPHOGENESIS_OF_AN_EPITHELIUM
Normalized Enrichment Score
B
Stemness-related pathways
DCLK1_MARUNO_ET_AL
GOBP_EPITHELIUM_DEVELOPMENT
GOBP_RESPONSE_TO_RETINOIC_ACID
GOBP_STEM_CELL_DIFFERENTIATION
IVANOVA_HEMATOPOIESIS_STEM_CELL_LONG_TERM
WP_WNT_SIGNALING_PATHWAY_AND_PLURIPOTENCY
GOBP_EPITHELIAL_TUBE_MORPHOGENESIS
GOBP_EMBRYONIC_MORPHOGENESIS
LIM_MAMMARY_STEM_CELL_UP
Normalized Enrichment Score
C
Immune system-related pathways
HALLMARK_INFLAMMATORY_RESPONSE
HALLMARK_INTERPHERON_GAMMA_RESPONSE
CARONNI_ET_AL_T1RS
GOBP_NEGATIVE_REGULATION_OF_LYMPHOCYTE_MEDIATED_IMMUNITY
GOBP_REGULATION_OF_LYMPHOCYTE_MEDIATED_IMMUNITY
GOBP_NEGATIVE_REGULATION_OF_LEUKOCYTE_PROLIFERATION
GOBP_ANTIMICROBIAL_HUMORAL_RESPONSE
GOBP_DEFENSE_RESPONSE_TO_BACTERIUM
HALLMARK_INTERPHERON_ALPHA_RESPONSE
GOBP_INTERFERON_MEDIATED_SIGNALING_PATHWAY
Normalized Enrichment Score
D
ECM pathways
REACTOME_ASSEMBLY_OF_COLLAGEN_FIBRILS_AND_OTHER_MULTIMERIC_STRUCTURES
REACTOME_DEGRADATION_OF_THE_EXTRACELLULAR_MATRIX
GOBP_FIBROBLAST_GROWTH_FACTOR_RECEPTOR_SIGNALING_PATHWAY
REACTOME_COLLAGEN_DEGRADATION
REACTOME_EXTRACELLULAR_MATRIX_ORGANIZATION
GOBP_RESPONSE_TO_FIBROBLAST_GROWTH_FACTOR
GOBP_CELL_CELL_JUNCTION_ORGANIZATION
Normalized Enrichment Score
E
Metabolic processes
GOBP_RESPONSE_TO_NUTRIENT
GOBP_RESPONSE_TO_VITAMIN
GOBP_REGULATION_OF_REACTIVE_OXYGEN_SPECIES_METABOLIC_PROCESS
Normalized Enrichment Score
F
tSNE_1
tSNE_2
Condition
CTRL
HFD
G
7.88% (N=199)
92.12% (N=2,327)
Hypermethylated
Hypomethylated
H
6.14% (N=155)
33.61% (N=849)
42.91% (N=1,084)
17.34% (N=438)
3'UTR
5' regulatory region
Body
Intergenic
I
Methylation
Condition
CTRL
HFD
J
Wnt signaling pathway
Cadherin signaling pathway
Count
Adjusted.P.value
Gene Ratio

***Figure S2. Murine GSEA and methylation analysis. A)*** *Gene Set Enrichment Analysis (GSEA) plot showing the enrichment in pathways associated with EMT and migration.* ***B****) GSEA plot showing the enrichment in pathways associated with stemness.* ***C)*** *GSEA plot showing the enrichment in pathways associated with the immune system and inflammatory processes.* ***D)*** *GSEA plot showing the enrichment in pathways associated with extracellular matrix (ECM).* ***E)*** *GSEA plot showing the enrichment in pathways associated with metabolic processes.* ***A-B-C-D-E)*** *FDR<0.25.* ***F)*** *t-SNE representing unsupervised analysis of control (CTRL) and high-fat diet (HFD) primary tumor cell lines for methylation assessment.* ***G-H)*** *Pie charts showing the number of differentially hyper and hypomethylated CpG sites between control and HFD cell lines (top panel) and the location of the sites (bottom panel).* ***I)*** *Heatmap showing the clustering of CTRL and HFD cell lines based on the differentially methylated CpG sites.* ***J)*** *Enrichment plot showing the pathways differentially methylated for HFD versus CTRL cell lines.*

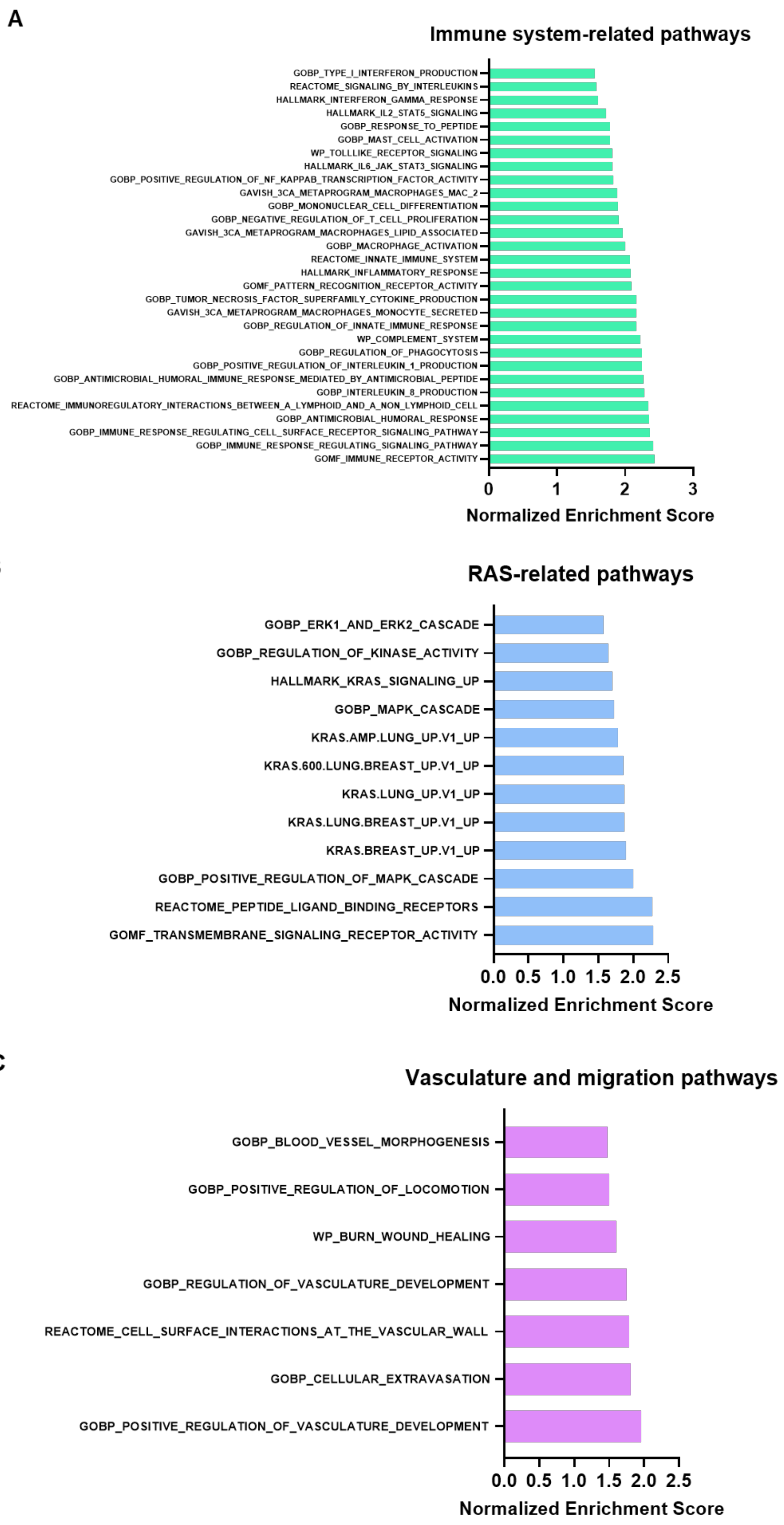
A
Immune system-related pathways
GOBP_TYPE_I_INTERFERON_PRODUCTION
REACTOME_SIGNALING_BY_INTERLEUKINS
HALLMARK_INTERFERON_GAMMA_RESPONSE
HALLMARK_IL2_STAT5_SIGNALING
GOBP_RESPONSE_TO_PEPTIDE
GOBP_MAST_CELL_ACTIVATION
WP_TOLLLIKE_RECEPTOR_SIGNALING
HALLMARK_IL6_JAK_STAT3_SIGNALING
GOBP_POSITIVE_REGULATION_OF_NF_KAPPAB_TRANSCRIPTION_FACTOR_ACTIVITY
GAVISH_3CA_METAPROGRAM_MACROPHAGES_MAC_2
GOBP_MONONUCLEAR_CELL_DIFFERENTIATION
GOBP_NEGATIVE_REGULATION_OF_T_CELL_PROLIFERATION
GAVISH_3CA_METAPROGRAM_MACROPHAGES_LIPID_ASSOCIATED
GOBP_MACROPHAGE_ACTIVATION
REACTOME_INNATE_IMMUNE_SYSTEM
HALLMARK_INFLAMMATORY_RESPONSE
GOMF_PATTERN_RECOGNITION_RECEPTOR_ACTIVITY
GOBP_TUMOR_NECROSIS_FACTOR_SUPERFAMILY_CYTOKINE_PRODUCTION
GAVISH_3CA_METAPROGRAM_MACROPHAGES_MONOCYTE_SECRETED
GOBP_REGULATION_OF_INNATE_IMMUNE_RESPONSE
WP_COMPLEMENT_SYSTEM
GOBP_REGULATION_OF_PHAGOCYTOSIS
GOBP_POSITIVE_REGULATION_OF_INTERLEUKIN_1_PRODUCTION
GOBP_ANTIMICROBIAL_HUMORAL_IMMUNE_RESPONSE_MEDIATED_BY_ANTIMICROBIAL_PEPTIDE
GOBP_INTERLEUKIN_8_PRODUCTION
REACTOME_IMMUNOREGULATORY_INTERACTIONS_BETWEEN_A_LYMPHOID_AND_A_NON_LYMPHOID_CELL
GOBP_ANTIMICROBIAL_HUMORAL_RESPONSE
GOBP_IMMUNE_RESPONSE_REGULATING_CELL_SURFACE_RECEPTOR_SIGNALING_PATHWAY
GOBP_IMMUNE_RESPONSE_REGULATING_SIGNALING_PATHWAY
GOMF_IMMUNE_RECEPTOR_ACTIVITY
0
1
2
3
Normalized Enrichment Score
B
RAS-related pathways
GOBP_ERK1_AND_ERK2_CASCADE
GOBP_REGULATION_OF_KINASE_ACTIVITY
HALLMARK_KRAS_SIGNALING_UP
GOBP_MAPK_CASCADE
KRAS.AMP.LUNG_UP.V1_UP
KRAS.600.LUNG.BREAST_UP.V1_UP
KRAS.LUNG_UP.V1_UP
KRAS.LUNG.BREAST_UP.V1_UP
KRAS.BREAST_UP.V1_UP
GOBP_POSITIVE_REGULATION_OF_MAPK_CASCADE
REACTOME_PEPTIDE_LIGAND_BINDING_RECEPTORS
GOMF_TRANSMEMBRANE_SIGNALING_RECEPTOR_ACTIVITY
0.0
0.5
1.0
1.5
2.0
2.5
Normalized Enrichment Score
C
Vasculature and migration pathways
GOBP_BLOOD_VESSEL_MORPHOGENESIS
GOBP_POSITIVE_REGULATION_OF_LOCOMOTION
WP_BURN_WOUND_HEALING
GOBP_REGULATION_OF_VASCULATURE_DEVELOPMENT
REACTOME_CELL_SURFACE_INTERACTIONS_AT_THE_VASCULAR_WALL
GOBP_CELLULAR_EXTRAVASATION
GOBP_POSITIVE_REGULATION_OF_VASCULATURE_DEVELOPMENT
0.0
0.5
1.0
1.5
2.0
2.5
Normalized Enrichment Score

***Figure S3. GSEA from human samples. A)*** *Gene Set Enrichment Analysis (GSEA) plot showing the enrichment in pathways associated with the immune system and inflammatory processes.* ***B)*** *GSEA plot showing the enrichment in pathways associated with RAS-related pathways.* ***C)*** *GSEA plot showing the enrichment in pathways associated with vasculature and migration. FDR<0.25.*

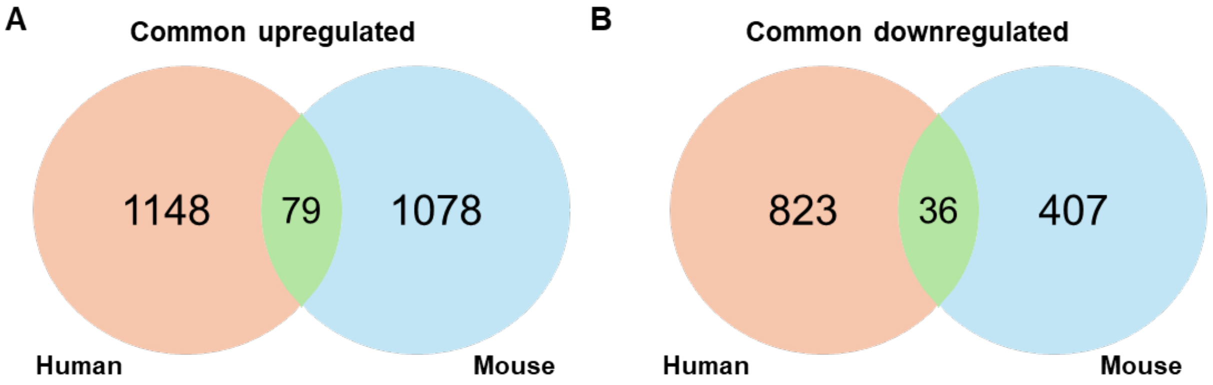


***Figure S4. Venn Diagrams of common up and downregulated genes in human and murine datasets. A-B)*** *Venn diagrams showing the number of genes commonly upregulated (A) or downregulated (B) found in our murine and human databases.*

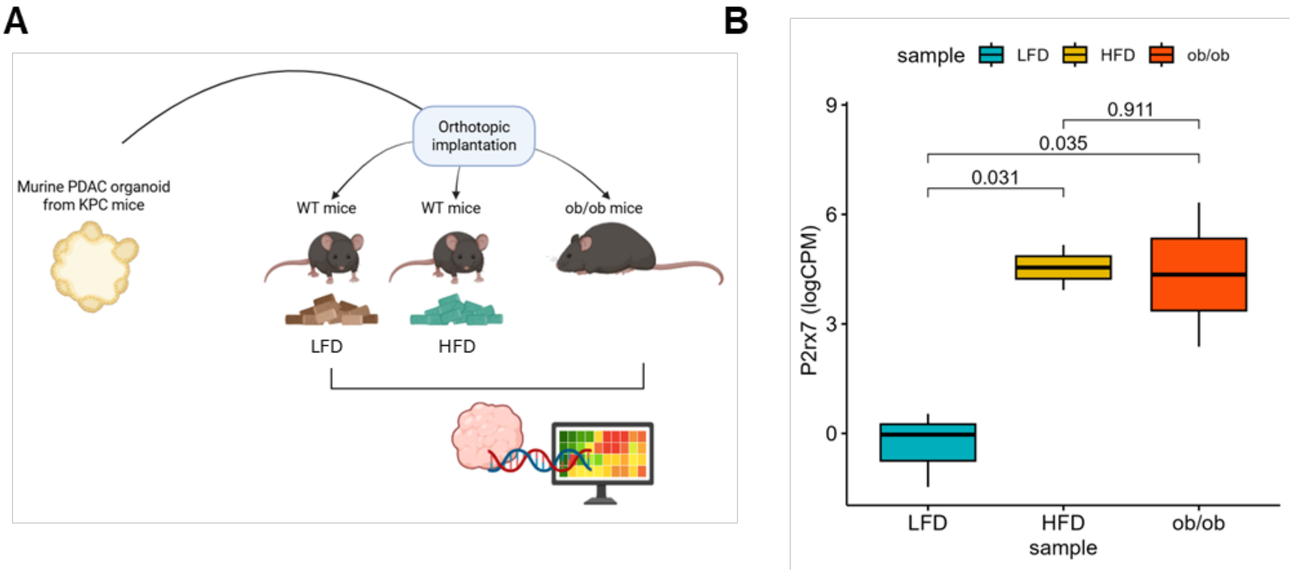


***Figure S5. Tumor-derived organoids grown in high-fat diet-fed and ob/ob mice recapitulate P2RX7 overexpression. A)*** *Schematic representation of the experimental setting to study how murine tumor-derived organoids generate tumors in three different hosts (C57Bl/6 mice fed with control or low fat (LFD) diet, HFD or ob/ob mice).* ***B)*** *Box plots showing the levels of P2rx7 expression by RNA-seq in tumors generated in the three different hosts, showing the mean ± STDEV of logCPM (n for LFD=3, n for HFD=2, n for ob7ob=2 P values as determined by Dunn test).*

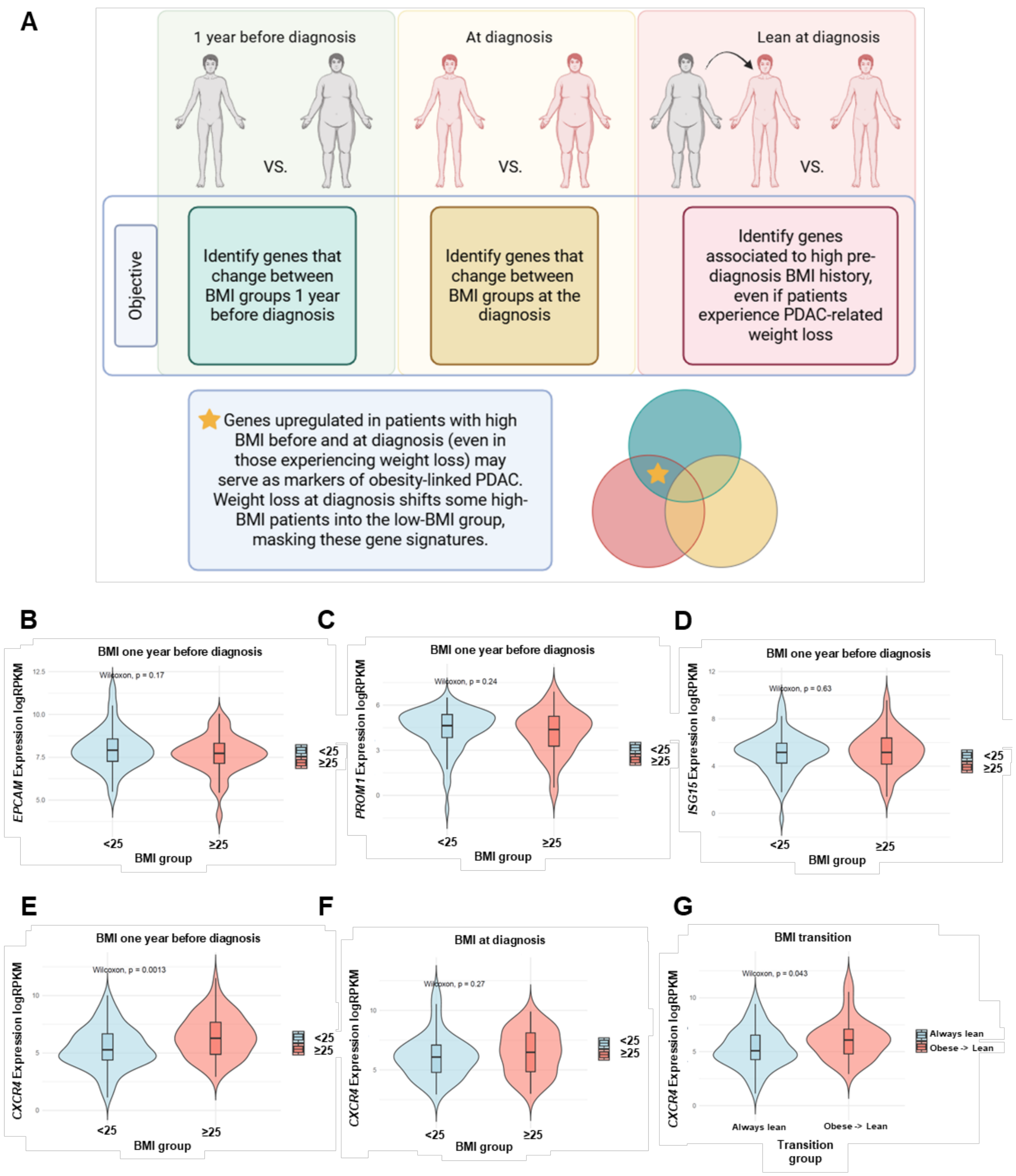


***Figure S6. Description of the comparisons in our patient cohort and evaluation of stem markers in patients with high and low BMI. A)** Diagram showing the comparisons employed to study the transcription diferences between patients with high BMI (BMI≥25) or low BMI (BMI<25) at different timepoints. **B)** Violin plot showing the quantification of EPCAM expression in PDAC patients based on their BMI 1 year before diagnosis. **C)** Violin plot showing the quantification of PROM1 expression in PDAC patients based on their BMI 1 year before diagnosis. **D)** Violin plot showing the quantification of ISG15 expression in PDAC patients based on their BMI 1 year before diagnosis. **E)** Violin plot showing the quantification of CXCR4 expression in PDAC patients based on their BMI 1 year before diagnosis. **F)** Violin plot showing the quantification of CXCR4 expression in PDAC patients based on their BMI at time of diagnosis. **G)** Violin plot showing the quantification of CXCR4*

*expression in PDAC patients based on their BMI transition (patients previously obese that lost their weight versus those that always present a BMI<25). P values as determined by Wilcoxon test.*

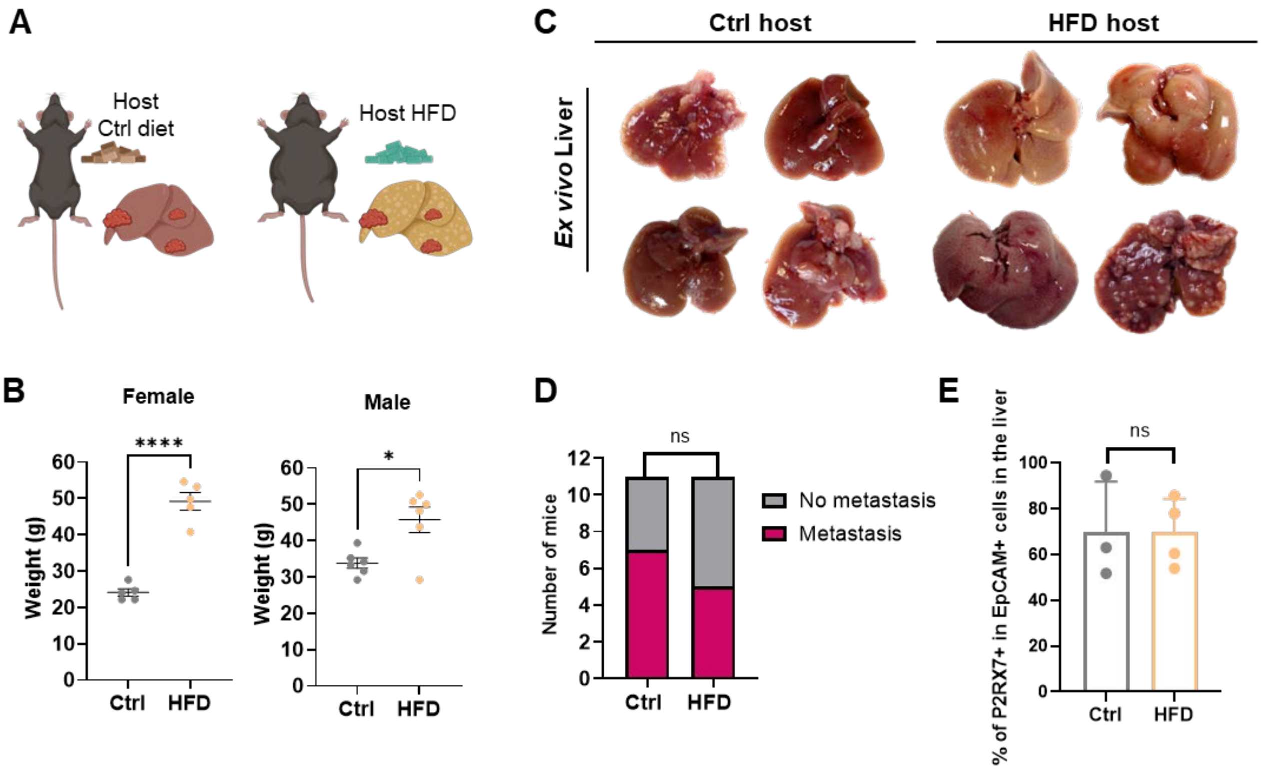


***Figure S7. Host-dependent metastatic capacity of control cells. A)*** *Diagram showing the two kinds of hosts employed for the intrasplenic assay with control (Ctrl) cell lines.* ***B)*** *Evaluation of body weight in female and male mice included in the study prior to the experiment. Shown is the mean ± SEM, P values as determined by unpaired T-test).* ***C)*** *Representative images of ex vivo livers from control and high-fat diet (HFD) mice with different degrees of macrometastatic disease.* ***D)*** *Number of mice presenting metastatic disease or not visible metastasis in the lean/control and HFD groups (n=11, P values as determined by Chi-square and Fisher´s exact test).* ***E)*** *Quantification of the percentage of P2RX7 positive cells in the detected EpCAM positive events in control and HFD livers from this experiment. Shown is the mean ± STDEV (n=3 mice, P values as determined by unpaired T-test). P values: *, <0.05; **, <0.01; ***, <0.001; ****, <0.0001.*

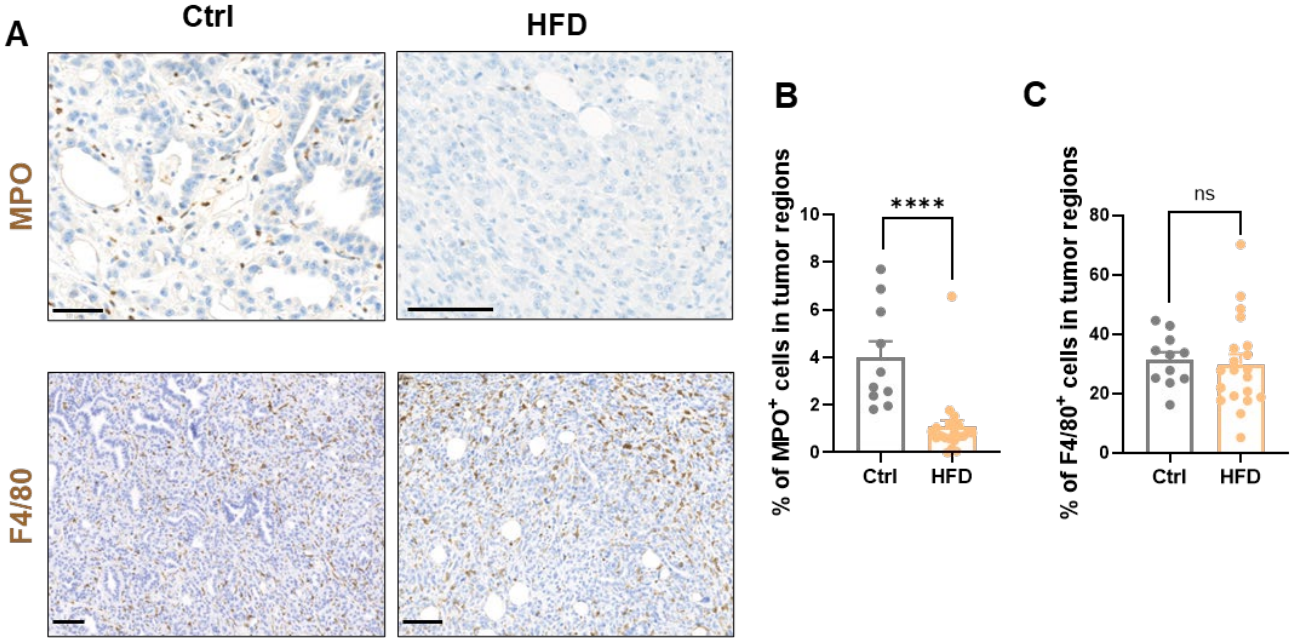


***Figure S8. Neutrophil and macrophage infiltration in HFD tumors. A)*** *Representative brightfield images of Ctrl and HFD mice tumors stained with anti-MPO (neutrophils) (scale bar = 50 µm on the left image and 100 µm on the right) or anti-F4/80 (macrophages) (scale bar = 100 µm) by immunohistochemistry.* ***B-C)*** *Quantification of MPO and F4/80 positive cells in pancreatic tumor regions in Ctrl and HFD mice. Shown is the mean ± SEM (n=3 mice for Ctrl group and 5 mice for HFD group, each dot represents a tumor region, P values as determined by unpaired T-test). P values: *, <0.05; **, <0.01; ***, <0.001; ****, <0.0001.*

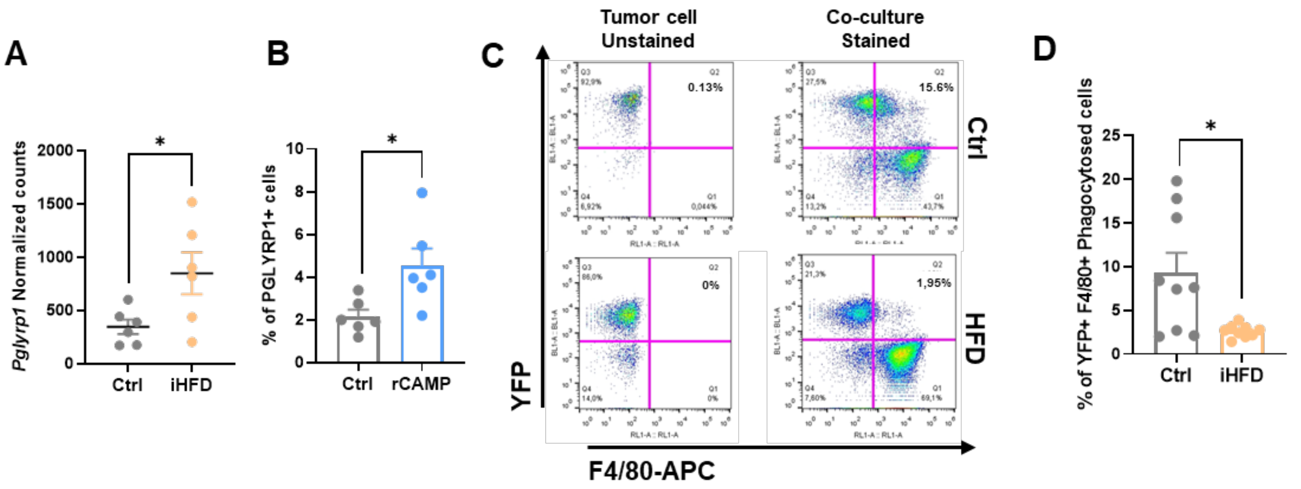


***Figure S9. CAMP induces P2RX7 and PGLYRP1 protecting tumor cells from macrophage phagocytosis. A)*** *Quantification of Pglyrp1 expression in control (Ctrl) and high-fat diet (HFD) primary murine cell lines included in the RNA-seq analyses. Shown is the mean ± SEM (n=6, P values as determined by T-test).* ***B)*** *Quantification of PGLYRP1 positive cells treated with recombinant (r) CAMP (100 µg/mL) for 48 hours. Shown is the mean ± SEM (n=6, P values as determined by unpaired T-test).* ***C)*** *Representative flow cytometry plots showing YFP positive control (Ctrl) and high-fat diet (HFD) primary tumor cells co-cultured with primary bone marrow-derived macrophages stained with F4/80. Live double positive events count as phagocytosed cells.* ***D)*** *Quantification of the percentage of phagocytosed tumor cells by flow cytometry. Shown is the mean ± SEM (n=3 cell lines in triplicates, P values as determined by unpaired T-test). P values: *, <0.05; **, <0.01; ***, <0.001; ****, <0.0001.*

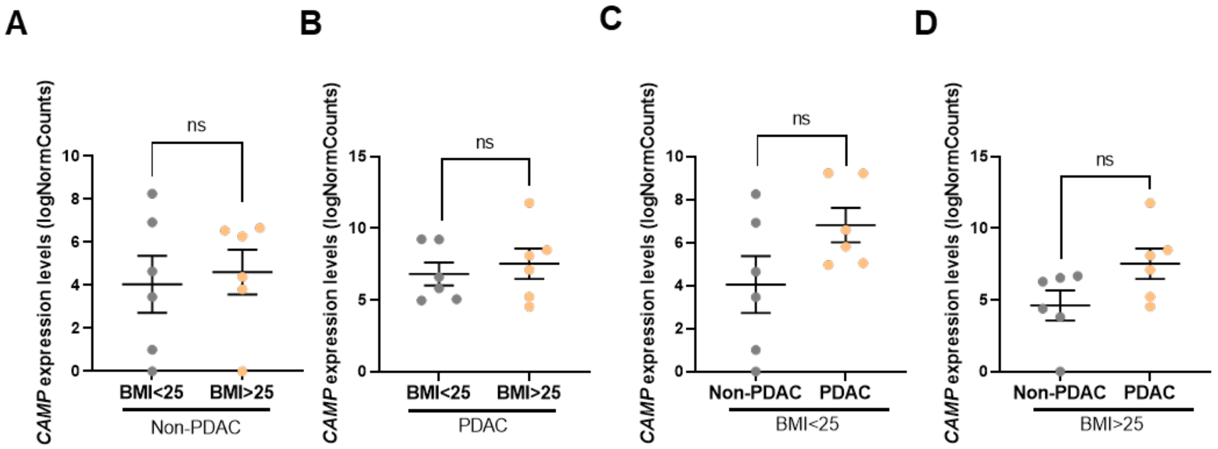


***Figure S10. CAMP can be detected in PBMCs from patients although it is not differentially expressed in circulation. A-B)*** *Quantification of CAMP expression levels in PBMCs from non-PDAC (A) and PDAC patients (B) with BMI higher or lower than 25. Shown is the mean ± SEM (n=6 patients per group, P values as determined by unpaired T-test).* ***C-D)*** *Quantification of CAMP expression levels in PBMCs from low BMI (C) and high BMI patients (D) either with non-PDAC or PDAC. Shown is the mean ± SEM (n=6 patients per group, P values as determined by unpaired T-test). P values: *, <0.05; **, <0.01; ***, <0.001; ****, <0.0001.*

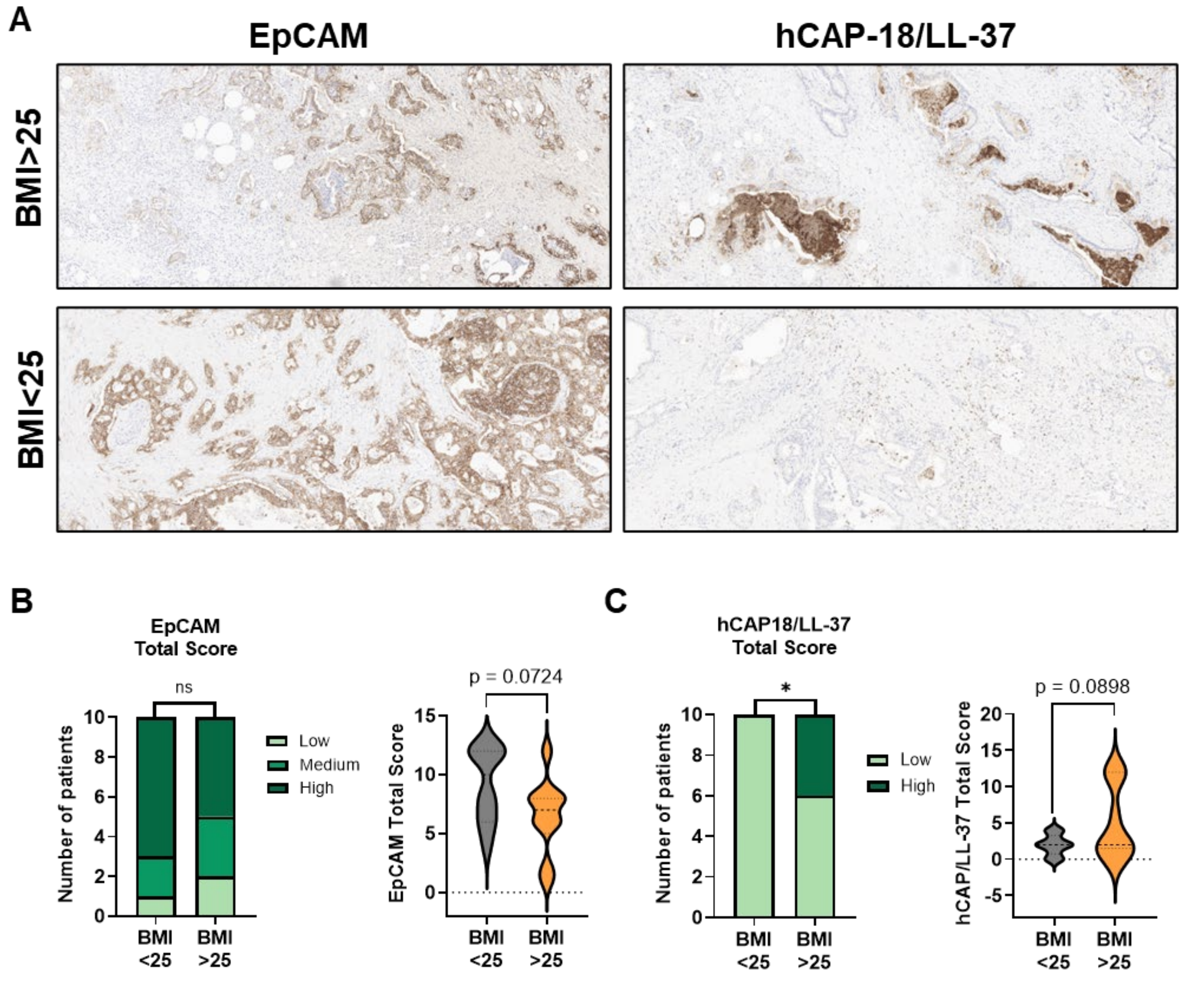

***Figure S11. Patients with higher BMI present lower EpCAM and higher hCAP-18/LL-37 intratumoral levels. A)*** *Representative brightfield images of PDAC tumors from patients with BMI>25 or BMI<25 stained with anti-EpCAM (epithelial tumor cells) or anti-hCAP-18/LL-37 by immunohistochemistry. Magnification for all images was 20x.* ***B)*** *Quantification of patient's proportion presenting high, medium or low EpCAM levels of the above mentioned stainings (left panel) and the EpCAM total score of all samples (right panel). Shown is the total score (% of staining x intensity), P values as determined by Chi-square and Fisher´s exact test (left panel) and unpaired T-test (right panel).* ***C)*** *Quantification of patient's proportion presenting high, medium or low hCAP-18/LL-37 levels of the above mentioned stainings (left panel) and the hCAP-18/LL-37 total score of all samples (right panel). Shown is the total score (% of staining x intensity), P values as determined by Chi-square and Fisher´s exact test (left panel) and unpaired T-test (right panel). P values: *, <0.05; **, <0.01; ***, <0.001; ****, <0.0001.*

### SUPPLEMENTARY TABLES

#### Supplementary Table 1

| Antibodies | | | | | |
|---|---|---|---|---|---|
| **Epitope** | **Host** | **Reactivity** | **Dilution** | **Application** | **Manufacturer (Cat no).** |
| CD133-APC | Rat | Mouse | 1:100 | Flow cytometry | eBioscience (17-1331-81) |
| EpCAM-FITC | Rat | Mouse | 1:100 | Flow cytometry | Miltenyi (130-123-674) |
| EpCAM-PE | Rat | Mouse | 1:100 | Flow cytometry | Miltenyi (130-117-779) |
| F4/80-APC | Rat | Mouse | 1:100 | Flow cytometry | Miltenyi (130-102-379) |
| PGLYRP1-AF555 | Mouse | Mouse/ Human | 1:100 | Flow cytometry | Novus Bio (NB100-56719AF555) |
| Sca-1-PE-Cy7 | Rat | Mouse | 1:100 | Flow cytometry | Biolegend (108124) |
| P2RX7 | Rabbit | Mouse/ Human | 1:50 | Flow cytometry | Sigma (P8997) |
| P2RX7-PE | Human | Mouse | 1:50 | Flow cytometry | Miltenyi (130-114-329) |
| CAMP | Rabbit | Mouse | 1:100 | Flow cytometry | Innovagen (PA-CRPL-10) |
| Secondary AlexaFluor 488 | Donkey | Rabbit | 1:200 | Flow cytometry | Invitrogen (A-21206) |
| CXCR4-APC | Human | Mouse | 1:100 | Flow cytometry | Miltenyi (130-123-274) |
| EpCAM | Rabbit | Mouse | 1:200 | IF | Abcam (ab71916) |
| Secondary Alexa Fluor 555 | Donkey | Rabbit | 1:200 | IF | Invitrogen (A31572) |
| Secondary Alexa Fluor 647 | Donkey | Mouse | 1:200 | IF | Invitrogen (A31571) |
| Vimentin | Mouse | Mouse | 1:100 | IF | Invitrogen (MA5-11883) |
| Cytokeratin 19 | Rat | Mouse | 1:750 | IHC | MONOCLONAL ANTIBODIES CORE UNIT |
| Vimentin | Rabbit | Mouse | 1:750 | IHC | Cell Signalling Technologies (5741) |
| F4/80 | Rabbit | Mouse | 1:750 | IHC | Bethyl (A700-209) |
| MPO | Rabbit | Mouse | 1:750 | IHC | DAKO (A 0398) |
| EpCAM | Mouse | Human | 1:400 | IHC | Cell Signalling Technologies (2929) |
| hCAP-18/LL-37 | Rabbit | Human | 1:750 | IHC | Innovagen (PA-LL37-100) |

#### Supplementary Table 2

| Gene | Species | Sequence Forward | Sequence Reverse |
|---|---|---|---|
| *ACTB* | Human | GCGAGCACACGAGCCTCGCCTT | CATCATCCATGGTGAGCTGGCGG |
| *Actb* | Mouse | CGGTTCCGATGCCCTGAGCTCTT | CGTCACACTTCATGATGGAATTGA |
| *Cdh1* | Mouse | TTCTGATCCTGCTGCTCCTACTG | TCTTCTTCTCCACCTCCTTCTTCATC |
| *Cdh2* | Mouse | CCATCATCGCTATCCTTCTGTGTATC | CGCTCTTTATCCCGCCGTTTC |
| *Krt19* | Mouse | AATGGCGAGCTGGAGGTGAAGA | CTTGGAGTTGTCAATGGTGGCAC |
| *Gata2* | Mouse | CTTCAACCATCTCGACTCGCAG | GCAACAAGTGTGGTCGGCACAT |
| *Tff2* | Mouse | GCATCACCAGTGAGCAGTGCTT | CTTGCGAGCTGACACTTCCATG |
| *Gata6* | Mouse | ATGCGGTCTCTACAGCAAGATGA | CGCCATAAGGTAGTGGTTGTGG |
| *P2rx7* | Mouse | GAACACGGATGAGTCCTTCGTC | CAGTGCCGAAAACCAGGATGTC |
| *P2RX7* | Human | CGACTAGGAGACATCTTCCGAG | GCAGTGATGGAACCAACGGTCT |
| *Camp* | Mouse | CTTCAACCAGCAGTCCCTAGAC | GCCACATACAGTCTCCTTCACTC |
| *Cxcr4* | Mouse | GACTGGCATAGTCGGCAATGGA | CAAAGAGGAGGTCAGCCACTGA |
| *Epcam* | Mouse | GAGTCCGAAGAACCGACAAGGA | GATGTGAACGCCTCTTGAAGCG |

#### Supplementary Table 3

| Name | Reference |
|---|---|
| Moffitt_2015_Classical_Top_100_MM | PMID: 26343385 |
| Moffitt_2015_Basal_Top_100_MM | PMID: 26343385 |
| MARTINELLI_GATA6_UP | PMID: 27325420 |
| MARTINELLI_GATA6_DOWN | PMID: 27325420 |
| MMU_PDAC_CSC | PMID: 38754953 |
| Youssef_EMT_T1_Invasive | PMID: 39414946 |
| Youssef_EMT_T2_Inflammatory | PMID: 39414946 |
| Youssef_EMT_Common | PMID: 39414946 |
